# Bacterial interspecies interactions modulate pH-mediated antibiotic tolerance in a model gut microbiota

**DOI:** 10.1101/538132

**Authors:** Andrés Aranda-Díaz, Benjamin Obadia, Tani Thomsen, Zachary F. Hallberg, Zehra Tüzün Güvener, Kerwyn Casey Huang, William B. Ludington

## Abstract

Despite decades of investigation into how antibiotics affect isolated bacteria, it remains highly challenging to predict consequences for communities in complex environments such as the human intestine. Interspecies interactions can impact antibiotic activity through alterations to the extracellular environment that change bacterial physiology. By measuring key metabolites and environmental pH, we determined that metabolic cross-feeding among members of the fruit fly gut microbiota drives changes in antibiotic sensitivity *in vitro*. Co-culturing of *Lactobacillus plantarum* with *Acetobacter* species induced tolerance to rifampin. Mechanistically, we found that acetobacters counter the acidification driven by *L. plantarum* production of lactate, and that pH shifts during stationary phase were sufficient to drive rifampin tolerance in *L. plantarum* monocultures. The key *Lactobacillus* physiological parameter related to tolerance was a reduction in lag time exiting stationary phase, opposite to a previously identified mode of tolerance to ampicillin in *E. coli. Lactobacillus* tolerance to erythromycin also depended on growth status and pH, suggesting that our findings generalize to other antibiotics. Finally, tolerance of *L. plantarum* to rifampin varied spatially across the fruit fly gut. This mechanistic understanding of the coupling among interspecies interactions, environmental pH, and antibiotic tolerance enables future predictions of growth and the effects of antibiotics in more complex communities and within hosts.

## Introduction

Decades of investigations have described detailed and precise molecular mechanisms of antibiotic action in model organisms. Yet, our current understanding is biased by a narrow set of standardized laboratory conditions; a recent study reported that resistance of *Escherichia coli* to the beta-lactam mecillinam is rarer in clinical isolates than in the laboratory and involves distinct genetic loci^1^. Unlike in laboratory monocultures, the vast majority of bacteria live in diverse communities such as the human gut microbiota. Antibiotics impact gut communities in many ways, ranging from the loss of diversity^2,3^ to the evolution of multidrug-resistant gut pathogens^4^. Hence, there is a pressing need for new frameworks that predict how antibiotics affect bacterial communities.

Bacteria can survive antibiotics through (i) resistance mutations, which counteract the antibiotic mechanism; (ii) persistence, whereby a subset of the bacterial population survives the antibiotic by becoming metabolically dormant; or (iii) tolerance, whereby the entire population enters an altered physiological state that is not susceptible to the antibiotic^5^. Members of multispecies communities, such as biofilms and models of urinary tract infections, can display altered sensitivity to antibiotics^6–9^. A few studies have delved into the molecular mechanisms behind cross-species antibiotic protection and sensitization. For example, the exoproducts of *Pseudomonas aeruginosa* affect the survival of *Staphylococcus aureus* through changes in antibiotic uptake, cell-wall integrity, and intracellular ATP pools^10^. In synthetic communities, intracellular antibiotic degradation affords cross-species protection against chloramphenicol^11^. Additionally, metabolic dependencies within synthetic communities can lower the viability of bacteria when antibiotics eliminate providers of essential metabolites, leading to an apparent change in the minimum inhibitory concentration (MIC) of the dependent species^9^. However, we still lack understanding of how contextual metabolic interactions between bacteria affect the physiological processes targeted by antibiotics and the resulting balance between growth inhibition (bacteriostatic activity) and death (bactericidal activity).

Characterizing the impact of metabolic interactions on antibiotic susceptibility requires functional understanding of how bacterial species interact during normal growth. Interspecies interactions can occur through specific mechanisms within members of a community (e.g. cross-feeding or competition for specific resources), or through global environmental variables modified by bacterial activity. An example of the latter is pH, which has recently been shown to drive community dynamics in a highly defined laboratory system of decomposition bacteria^12^.

While synthetic communities afford the opportunity to design and to tune bacterial interactions, it is unclear whether findings are relevant to natural communities. The stably associated gut microbiota of *Drosophila melanogaster* fruit flies constitutes a naturally simple model community for determining how metabolic interactions between species affect growth, physiology, and the action of antibiotics^13^. This community consists of ~5 species predominantly from the *Lactobacillus* and *Acetobacter* genera^14^ (Fig. 1a). Lactobacilli produce lactic acid^15^, while acetobacters are acetic acid bacteria that are distinguished by their ability to oxidize lactate to carbon dioxide and water^16^. Short chain fatty acids, such as lactate, decrease the pH of natural fermentations and may constitute a mechanism through which pH plays a prominent role in community dynamics. The naturally low number of species in *Drosophila* gut microbiota and its compositional modularity (lactobacilli versus acetobacters) enable systematic dissection of microbial interactions.

**Figure 1:**
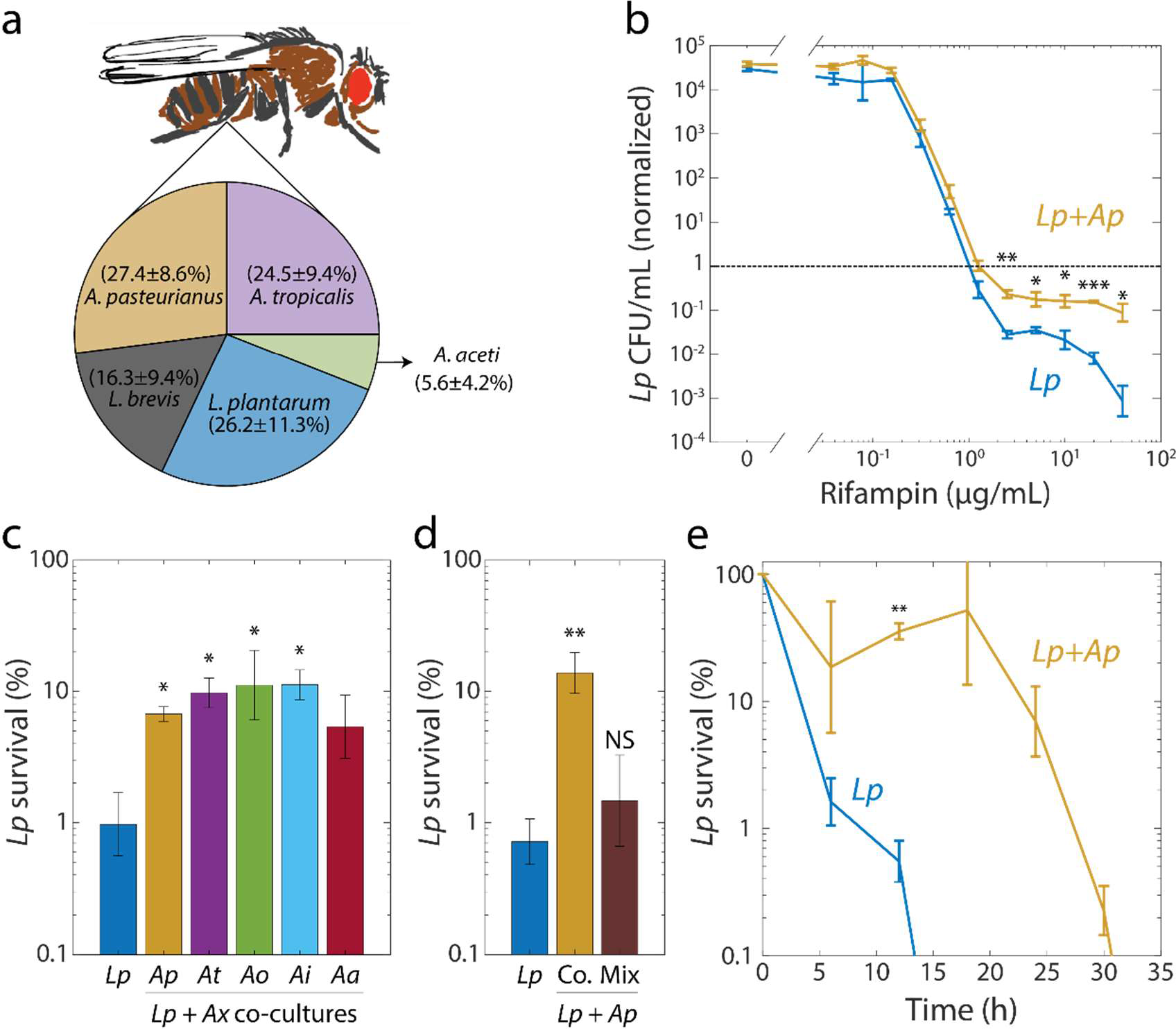
Interspecies interactions within the fruit fly gut microbiome induce rifampin tolerance. a) Relative abundances of the dominant species in the *D. melanogaster* gut microbiome determined from 16S rRNA sequencing. Values are mean ± standard deviation (S.D.), *n*=18. Mean and S.D. were weighed by the total number of reads for each fly. b) When grown with *Ap, Lp* survived after 24 h at rifampin concentrations above the MIC. Viable cell plating counts of *Lp* after growth in rifampin for 24 h normalized to the counts at the start of the experiment (*t*=0). Error bars are standard deviation (S.D.) for each condition, *n*=3. *P*-values are from a Student’s two-sided *t*-test of the difference of the co-culture from the monoculture (*: *P*<4×10^−3^, **: *P*<8×10^−3^, ***: *P*<8×10^−5^). c) Protection of *Lp* at supra-MIC concentrations of rifampin is elicited by all acetobacters tested. Normalized CFUs of *Lp* grown in monoculture (*Lp*) or in co-culture with *Ap, At, Ao, Ai*, and *Aa*, and then treated with 20 μg/mL rifampin for 24 h. Error bars are S.D. for each condition, *n*=3. *P*-values are from a Student’s two-sided *t*-test of the difference from the monoculture (*: *P*<0.01). d) *Ap*-mediated survival of *Lp* at rifampin concentrations above the MIC is history-dependent, requiring co-culturing before exposure as compared with mixing. Normalized CFUs of *Lp* grown in monoculture, in co-culture with *Ap* (Co.), or mixed with *Ap* without subsequent growth in the absence of antibiotic (mix), and treated with 20 μg/mL rifampin for 24 h. Error bars are S.D. for each condition, *n*=3. *P*-values are from a Student’s two-sided *t*-test of the difference from the monoculture (**: *P*<5×10^−3^, NS: not significant). e) The time to killing of *Lp* under rifampin treatment is extended in the presence of an Acetobacter. Normalized CFUs of *Lp* grown in monoculture and co-cultured with Ap, and treated with 50 μg/mL rifampin. Error bars are S.D. for each condition, *n*=3. *P*-values are from a Student’s two-sided *t*-test of the difference from the monoculture at the corresponding timepoint (**: *P*<1×10^−3^). Values off the graph were below the limit of detection of the assay

In the current study, we interrogated how interspecies interactions affect growth and antibiotic susceptibilities. We used high-throughput assays to measure these parameters in monocultures versus co-cultures and inside fly guts. *Lactobacillus plantarum* (*Lp*) exhibited antibiotic tolerance (delay in death^5^ by ~12 h) in the presence of acetobacters. Lactate accumulation by *Lp* in monocultures acidified the media, inhibiting growth during stationary phase. *Acetobacter*-mediated lactate consumption released this inhibition by increasing pH, leading to a shorter *Lp* lag while exiting stationary phase. This reduced lag exiting stationary phase corresponded with the antibiotic tolerance of *Lp* that we observed. We determined that changes in pH elicited by *Acetobacter* activity are sufficient to modulate tolerance of *Lp* to both rifampin and erythromycin. Finally, *ex vivo* experiments revealed that antibiotic tolerance differs in distinct compartments of the host gastrointestinal tract. Taken together, our findings indicate that simple changes to the environment can drive complex behaviors within bacterial communities.

## Results

### Interspecies interactions induce tolerance to rifampin

To determine the composition of the gut microbiota in our laboratory fruit flies, we performed deep sequencing of 16S rRNA V4 amplicons from 18 individual dissected guts (Methods). We identified five species belonging to seven unique operational taxonomic units (OTUs) by clustering the sequences at 99% identity: *L. plantarum* (*Lp*), *L. brevis* (*Lb*), *Acetobacter pasteurianus* (*Ap*), *A. tropicalis* (*At*), and *A. aceti* (*Aa*) (Fig. 1a). We then isolated the species in culture and determined the antibiotic sensitivities of the four major fly gut inhabitants (*Lp, Lb, Ap* and *At*; Fig. 1a) *in vitro* using isolates of these species (Table S1) grown in Man, Rogosa, and Sharpe (MRS) medium. We tested 10 antibiotics representing a wide variety of classes using plate-based growth assays (Methods). For many drugs, some of the fly gut species were resistant (detectable growth) at least up to the highest concentrations tested. Rifampin was the only drug for which all four species exhibited sensitivity (Table S2) and it is bactericidal^17^, hence there is the opportunity to study survival as well as sensitivity.

We noted from growth curves in the absence of drug that *Lb* grew significantly more slowly than *Lp* (0.60 ± 0.054 h^−1^ vs. 0.64 ± 0.001 h^−1^ for *Lp, P* = 5.5×10^−3^, *n* = 16, Fig. S1) and had a much longer lag phase than *Lp* (6.55 ± 0.16 h vs. 1.92 ± 0.08 h for *Lp, P* = 1.8×10^−39^, *n* = 16, Fig. S1). Thus, we focused on *Lp* and its interactions with the *Acetobacter* species, particularly *Ap*, which is more abundant in the fly gut than the other acetobacters (Fig. 1a).

We grew *Lp* and *Ap* separately for 48 h in test tubes, combined them in test tubes at an optical density at 600 nm (henceforth OD) of 0.02 each, and co-cultured them in MRS for 48 h. We then diluted this co-culture and 48-h monocultures of *Lp* and *Ap* into fresh MRS at ~5×10^5^ colony-forming units/mL (CFU/mL) in 96-well plates with various concentrations of rifampin and measured growth over 48 h. The MIC of the co-culture was similar to that of *Lp* alone (2.5 μg/mL, Fig. S2a). To determine whether the co-culture still contained both species, we measured the percentage of survival and the fraction of each species at various rifampin concentrations by taking advantage of the fact that *Lp* and the acetobacters have distinct colony morphologies and colors on MRS and MYPL plates (Table S1). We measured *Lp* CFUs on MRS and *Ap* CFUs on MYPL because *Lp* and *Ap* grow more quickly on MRS and MYPL, respectively. In the co-culture, *Ap* died off at a similar concentration of rifampin as during growth in a monoculture (1.25 μg/mL, Fig. S2b). For *Lp*, the MIC was the same in co-culture as in monoculture (1.25 μg/mL, Fig. 1b), but at concentrations above the MIC, significantly more *Lp* cells survived in co-culture than in monoculture (Fig. 1b). This effect could not be explained by small differences in the initial inoculum, as increasing cell densities up to 100-fold did not change the MIC or survival of *Lp* in monoculture (Fig. S2c,d). Because of the change in *Lp* survival, we focused herein on this phenotype.

To determine whether *Lp*’s increased survival was specific to co-culturing with *Ap*, we co-cultured *Lp* with each of the acetobacters, including a wild fly isolate of *A. indonesiensis* (*Ai*), and lab fly isolates of *A. orientalis* (*Ao*) and *Aa*, the fifth major component of the microbiota of our flies (Fig. 1a). We then diluted each co-culture to an initial *Lp* cell density of ~5×10^5^ CFU/mL into fresh MRS with 20 μg/mL rifampin (16X MIC) and let the cells grow for 24 h. Co-culturing with any of the acetobacters increased survival by approximately one order of magnitude (Fig. 1c). To determine whether this increased survival requires co-culturing prior to rifampin treatment (rather than the presence of the acetobacters being sufficient), we grew *Lp* and *Ap* separately for 48 h and mixed and diluted them at the time of addition of 20 μg/mL rifampin. After 24 h of growth, the number of CFU/mL was significantly lower in mixed culture than in co-culture (Fig. 1d), indicating a history dependence to increased survival.

To determine whether co-culturing slows killing by the drug, we examined the survival of *Lp* over time at a high drug concentration (50 μg/mL, 40X MIC). Similar to the experiments above, we compared *Lp* CFU/mL in a monoculture with that in a co-culture with *Ap*. In monoculture, *Lp* rapidly died, with CFU/mL becoming undetectable within 18 h; in contrast, *Lp* survived >30 h after co-culturing (Fig. 1e). This increased time to death of *Lp* as a bulk population, and unchanged MIC, together indicate that co-culturing *Lp* with *Ap* induces tolerance of *Lp* to rifampin^5^.

### *Co-culturing leads to growth of* Lp *in stationary phase*

Our finding that co-culturing *Lp* with acetobacters affects antibiotic tolerance (Fig. 1c,e) prompted us to investigate the environmental factors that cause this phenotype. We first inquired whether the total amount of growth of the co-culture was larger or smaller than expected from the yield of the monocultures. We grew *Lp* and each of the acetobacters separately for 48 h, diluted the monocultures to OD = 0.04, combined the *Lp* monoculture 1:1 with each Acetobacter monoculture, and grew the co-cultures for 48 h in test tubes. In bulk measurements, the *Lp-Ap* co-culture showed a significant synergistic effect (Supplementary Text, Fig. S3a). We then determined the total carrying capacity of each of the species in the co-cultures grown in test tubes by counting CFUs.

We determined that the *Lp* CFU/mL values for 48-h co-cultures with *Ap, At*, and *Ai* were higher than the *Lp* CFU/mL values in monoculture (Fig. 2a). Co-cultures with *Ap* showed the strongest effect; *Aa* and *Ao* did not significantly increase *Lp* CFU/mL (Fig. 2a). All acetobacters except for *Ap* reached lower CFU/mL in co-cultures with *Lp* than in monocultures (Fig. S3b). Thus, *Lp* has a strong positive interaction with *Ap*, and negative or neutral interactions with the rest of the acetobacters (Supplementary Text, Fig. S3c).

**Figure 2:**
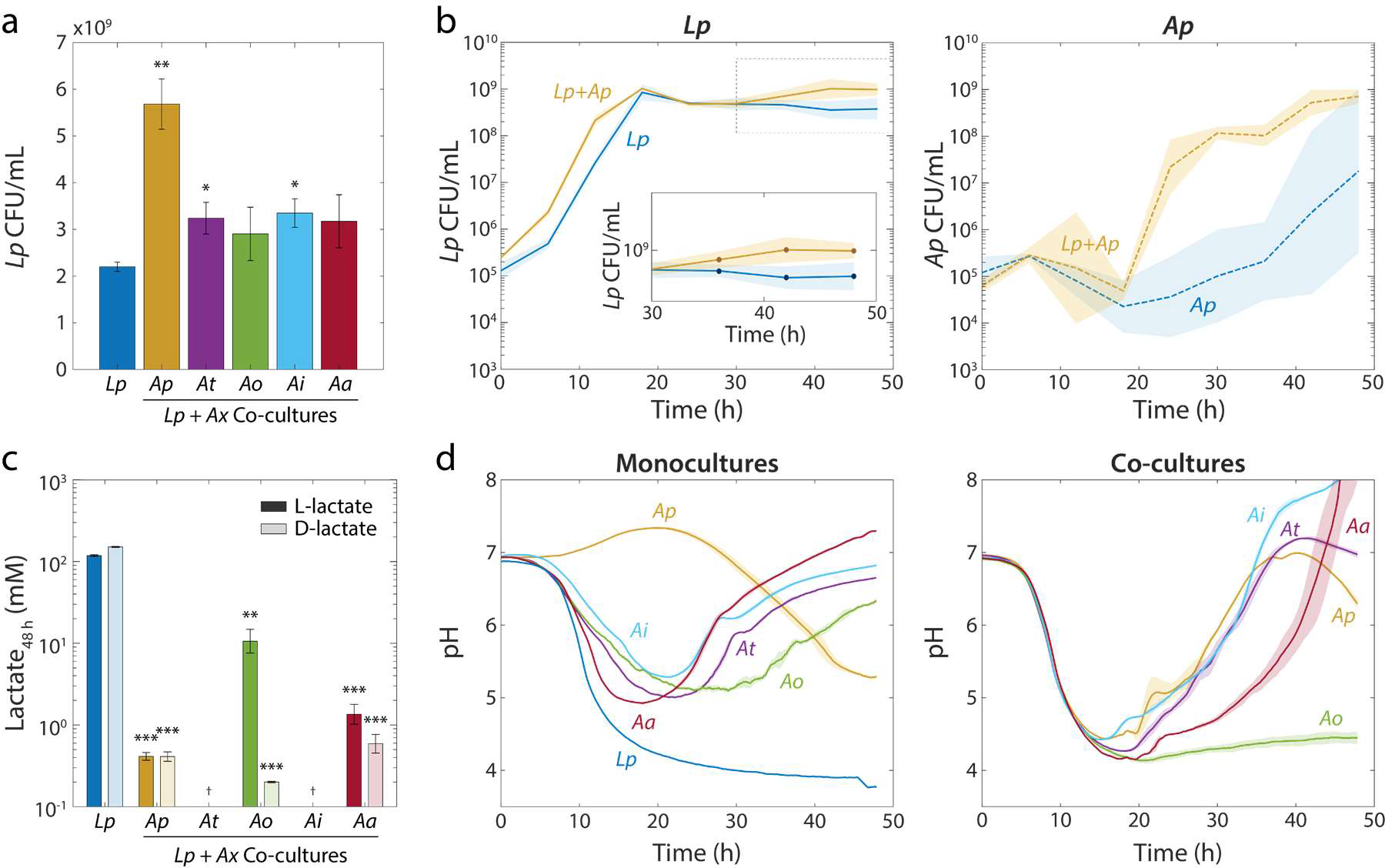
*Lp* growth during stationary phase in *Acetobacter* co-cultures is associated with an increase in pH and a decrease in lactate concentration. a) Co-culturing *Lp* with *Ap, At, Ai*, or Aa resulted in increased *Lp* cell density after 48 h. Co-culturing with *Ao* did not significantly increase *Lp* cell density by 48 h. Error bars are standard deviation (S.D.) for each condition, *n*=3. *P*-values are from a Student’s two-sided t-test of the difference from the monoculture (*: *P*<0.01, **: *P*<2×10^−3^). b) Co-culturing *Lp* with *Ap* resulted in higher *Lp* cell density in stationary phase, as well as faster growth and shorter lag for *Ap*. Shaded regions indicate S.D., *n*=3. Inset: zoom-in on region inside dashed box highlighting increase in carrying capacity in co-culture. c) L- and D-lactate were produced in *Lp* monocultures and consumed in co-cultures. Lactate concentration was measured enzymatically from culture supernatants at 48 h. Error bars are S.D. for each condition, *n*=3. *P*-values are from a Student’s two-sided t-test of the difference from the monoculture (**: *P*<2×10^−3^, ***: *P*<2×10^−4^). d) The increase in *Lp* cell density in stationary phase is associated with an Acetobacter-dependent increase in pH early in stationary phase. pH was measured with the pH-sensitive dye 2’,7-bis-(2-carboxyethyl)-5-(and-6)-carboxyfluorescein over time (Methods). Shaded regions indicate S.D., *n*=3.

To determine when the additional growth took place, we monitored CFU/mL values for *Lp* and *Ap* in co-culture throughout a 48-h time course starting from an initial combined cell density of ~5×10^5^ CFU/mL. Initially, *Lp* accounted for the bulk of the growth in the co-culture (Fig. 2b). Interestingly, *Ap* in liquid monoculture showed little to no growth in most replicates after 40 h (Fig. 2b); by contrast, in liquid co-culture *Ap* started to grow after ~20 h and reached saturation by ~40 h (Fig. 2b), indicating that *Ap* also benefited from growth as a co-culture. This benefit likely stemmed from a reduction in *Ap* cell death during lag phase (Supplementary Text, Fig. S4). Thus, a mutualism exists between *Lp* and *Ap* driven by growth during and exiting from stationary phase.

### Lactate metabolism leads to changes in pH in stationary phase co-cultures

Interestingly, after 30 h, *Lp* displayed a significant (~2X) increase in CFU/mL in the co-culture that did not occur in the monoculture (Fig. 2b), indicating that the increase in final yield occurs late in stationary phase. We therefore hypothesized that *Lp* has a common metabolic interaction with each of the acetobacters. An obvious candidate is cross-feeding, since *Lp* produces lactate and the acetobacters consume it. We measured lactate levels in the supernatants of *Lp* monocultures and co-cultures of *Lp* with each of the acetobacters individually, after 48 h of growth. As expected, the *Lp* monoculture accumulated L- and D-lactate to high levels (>100 mM; Fig. 2c). All co-cultures had significantly lower concentrations of both isomers than the monoculture (<2 mM, Fig. 2c). The *Lp-Ao* co-culture harbored higher levels of L-lactate than any other co-culture and *Lp-Aa* had higher concentration of L-lactate than the rest of the co-cultures (Fig. 2c). *Lp-Ap*, *Lp-Ao*, and *Lp-Aa* co-cultures all accumulated lactate (>10 mM) at 20 h (Fig. S5a). Taken together, these data suggest that *Lp* metabolism leads to an initial accumulation of lactate and that the acetobacters consume it, although *Aa* and *Ao* are less efficient at consuming L-lactate than the other species.

Since lactate is a short-chain fatty acid with a p*K*_a_ of 3.86, we suspected that lactate production would affect the pH of the culture. We first monitored the pH dynamics of monocultures of *Lp* and of each *Acetobacter* using the pH-sensitive fluorophore 2’,7-bis-(2-carboxyethyl)-5-(and-6)-carboxyfluorescein (BCECF)^18^. In the *Lp* monoculture, pH decreased from pH=6.75 to below 4 during growth (Fig. 2d); more precisely, we measured a final supernatant pH=3.77 using a pH meter (Fig. S5a). We measured pH over time in *Acetobacter* monocultures using BCECF. For all acetobacters except *Ap*, the medium first acidified down to pH~5, and then increased back to pH=6-7 (Fig. 2d).

To test whether acetobacters reverse the pH decrease due to the accumulation of lactate produced by *Lp*, we measured the pH of co-cultures over time using BCECF. Co-cultures with *Ap, At*, and *Ai* followed similar trajectories in which the pH followed that of the *Lp* monoculture for the first 20 h, after which the pH increased up to a final value of ~7 (Fig. 2f). The *Lp-Aa* co-culture experienced a ~10-h delay in the pH increase, while the co-culture with *Ao* showed only a slight pH increase by 48 h (Fig. 2f). The slight pH increase in *Ao* co-culture is consistent with lower L-lactate consumption by this species (Fig. 2c). Using a pH meter for validation, we measured final pH values of 5.9, 5.8, 4.6, 5.4, and 4.8 in co-cultures with *Ap, At, Ao, Ai*, and *Aa*, respectively (Fig. S5b). Thus, lactate metabolism dictates dramatic shifts in environmental pH that are related to physiological changes in antibiotic tolerance (Fig. 1c).

Given the strong acidification of the medium in *Lp* monoculture but not in co-culture (Fig. 2d), we hypothesized that intracellular pH decreases in monoculture and increases in co-culture. To measure intracellular pH, we transformed our *Lp* strain with a plasmid expressing pHluorin (a GFP variant that acts as a ratiometric pH sensor^19^) under the control of a strong constitutive promoter^20^. The two absorbance peaks, which we measured at 405 and 475 nm, are sensitive to pH and the ratio of the emission (at 509 nm) at these two excitation wavelengths can be used to estimate intracellular pH. We grew this strain in monoculture and in co-culture with *Ap* and measured fluorescence over time in a plate reader. Because of the high autofluorescence of the medium at 405 nm (data not shown), we could only track changes in fluorescence at an excitation wavelength of 475 nm. We observed an initial increase in signal as the *Lp* cells started to proliferate (Fig. S6a). After the cultures saturated (*t*~20 h), we detected a decrease in the signal down to the levels of medium autofluorescence in the monoculture (Fig. S6a). In the co-culture, where the extracellular pH is raised by the metabolic activity of *Ap*, fluorescence did not decrease over time (Fig. S6a), suggesting that intracellular pH decreases in a time-dependent manner in monoculture but not in co-culture.

To verify that the decrease in fluorescence in monoculture was due to a drop in intracellular pH, as opposed to a decrease in protein synthesis, we sampled cells after 48 h of growth, centrifuged them, resuspended them in PBS in order to measure pHluorin signal at both its excitation wavelengths, and measured fluorescence within 1 minute of resuspension. The ratio of the signal from pHluorin at its two excitation wavelengths was significantly higher in co-culture (Fig. S6b). Taken together, these data indicate that the intracellular pH of *Lp* cells is significantly lower in monoculture than in co-culture with acetobacters.

### *Low pH inhibits the growth of* Lp *and extends lag phase*

Since *Ap* growth causes a large increase in the extracellular pH of an *Lp-Ap* co-culture, we sought to determine the dependence of *Lp* growth on pH. We diluted a 48-h culture of *Lp* cells grown in MRS at starting pH=6.75, to a starting OD=0.02 in MRS adjusted to starting pH ranging from 3 to 8 (Methods). We then measured growth and BCECF fluorescence in a plate reader (Fig. S7a,b). For lower starting pH values, the carrying capacity was lower (Fig. 3a) and varied over a large OD range from <0.02 to >2. For all starting pH values, *Lp* cells reduced the pH to a common final value of ~3.7 (Fig. 3a). The bulk growth rate approached zero (Fig. 3b) as the pH approached its final value, explaining the differences in yield. Interestingly, the maximum growth rate was also pH-dependent (Fig. 3c), with the highest growth rate at starting pH=7.

**Figure 3:**
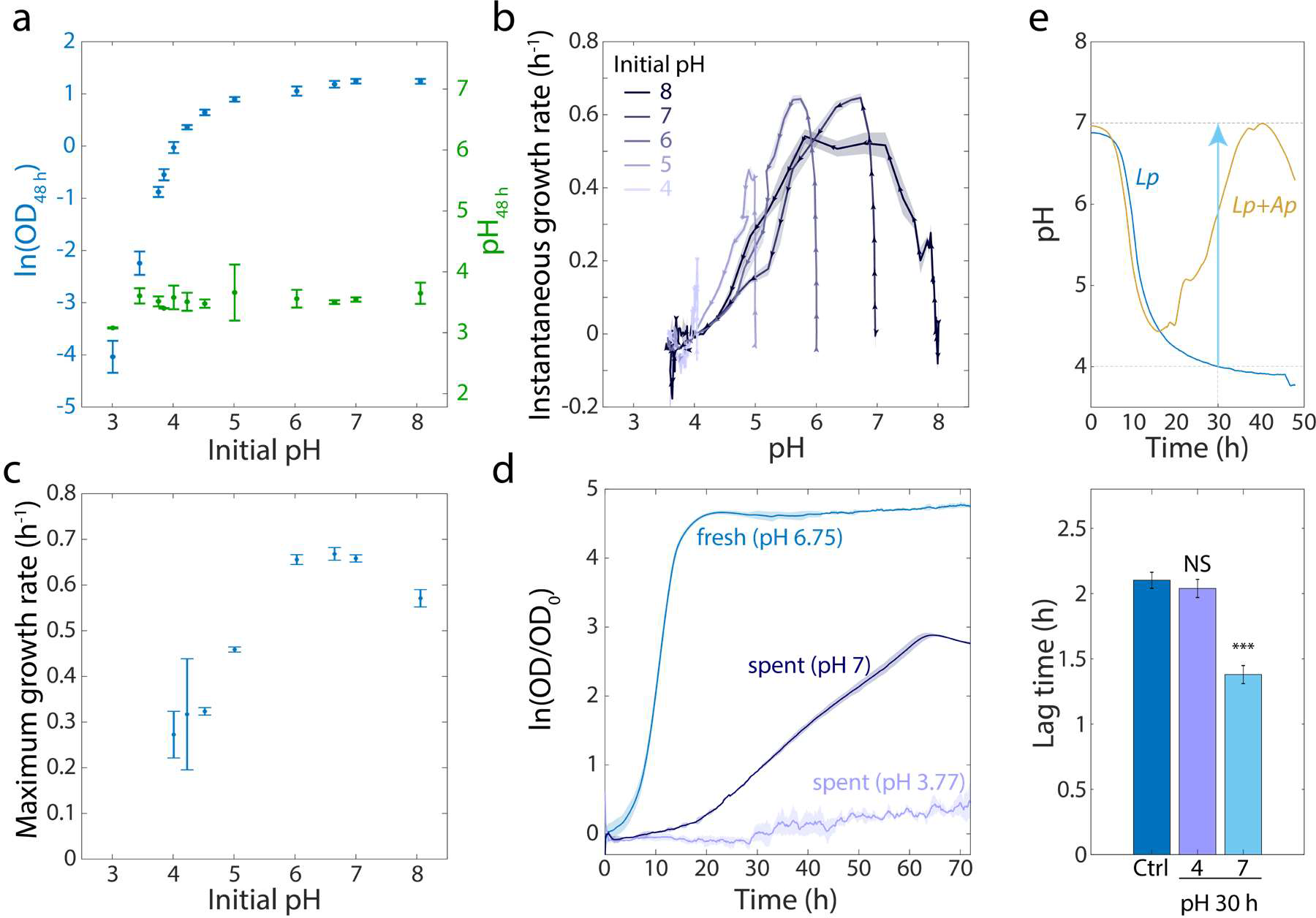
An increase in extracellular pH in stationary phase releases growth inhibition in *Lp* monocultures and shortens lag phase. a) *Lp* growth is inhibited by low pH. Logarithm of OD (blue) and pH measured using BCECF (green) after 48 h of growth in MRS at various starting pH values. Error bars are standard deviation (S.D.), *n*=4. b) Instantaneous growth rate in MRS is strongly linked to pH. Each curve was initialized at a different starting pH and represents 48 h of growth. Arrowheads indicate direction of time. Shaded regions are S.D., *n*=4. c) Maximal growth rate in MRS increases with increasing initial pH. Error bars are S.D., *n*=4. d) Increasing the pH of a saturated, spent *Lp* culture from 3.77 to 7 allows growth, although not as much as fresh MRS. Error bars are S.D., *n* = 3. e) Increasing the pH of an *Lp* monoculture at *t* = 30 h from 4 to 7 to mimic the pH increase in *Lp-Ap* co-culture (top) leads to a shorter lag phase (bottom). Lag time was calculated by fitting growth curves to the Gompertz equation. Error bars are S.D., *n*=3. *P*-values are from a Student’s two-sided t-test of the difference from the control (***: *P*<5×10^−4^, NS: not significant).

Given these findings, we hypothesized that the inhibition of growth in stationary phase of an *Lp* monoculture is due to the decreased intracellular and extracellular pH, and that *Ap* releases *Lp*’s growth inhibition by raising intracellular and extracellular pH. To test this hypothesis, we inoculated a 48-h culture of *Lp* to an initial OD=0.02 into the supernatant of a 48-h *Lp* culture at pH=3.77 or set to pH=7. We observed substantially more growth (~20-fold increase) of the bulk culture in supernatant raised to pH=7, while no growth took place starting from pH=3.77 (Fig. 3d). As expected, the maximal growth rate was lower than in fresh MRS due to the partial depletion of nutrients (Fig. 3d); addition of glucose to the conditioned medium supported faster growth, but only starting from neutral pH (Fig. S7c). We also hypothesized that the accumulation of lactate by *Lp* would allow growth of *Ap* in *Lp*-conditioned medium even at low starting pH. When we diluted a saturated *Ap* culture into *Lp*-conditioned medium generated as above, *Ap* grew to similar levels as in fresh MRS (Fig. S7d).

These findings suggested that the effects of *Ap* in co-culture on *Lp* growth might be due primarily to the pH changes that *Ap* initiates because of the ability of acetobacters to grow at low pH and to consume lactate. Thus, we first increased the pH of an *Lp* monoculture to 7 after 30 h, when the pH increased most rapidly in the *Lp-Ap* co-culture (Fig. 3e), and incubated cells for an additional 18 h. We did not observe a significant increase in CFU/mL from this pH-adjusted culture versus controls that simply grew for 48 h or were subjected to all washes required for pH adjustment and then returned to the same supernatant (Fig. S7e). We then assessed if increasing the pH at 30 h resulted in a decrease in the duration of lag phase by diluting the monoculture to OD=0.0375 into fresh MRS after an additional 18 h of growth. The lag phase was shorter in the pH-adjusted culture than control bulk cultures (Fig. 3e). Thus, pH is a driver of the growth advantages of *Lp* in lag phase.

### *Co-culturing* Lp *with acetobacters reduces lag time*

Canonical antibiotic tolerance in *E. coli* results from a decrease in growth rate or an increase in lag phase that protects cells through metabolic inactivity^5^. To measure growth rate and lag phase, we co-cultured *Lp* with each of the acetobacters individually for 48 h, diluted the culture to a common OD of 0.0375, and monitored growth in a plate reader. The maximum growth rate was the same for the *Lp* monoculture and co-cultures with *Ap, At*, and *Ai*, and slightly higher for co-cultures with *Ao* and *Aa* (Fig. S8). We previously observed for *Lp* monocultures when shifting the pH that the stimulation of growth in stationary phase was connected with tolerance (Fig. 3e), opposite to that of *E. coli* tolerance to ampicillin^5^. In agreement with these data, there was a significant decrease in bulk lag time for all of the Acetobacter co-cultures (Fig. 4a,b). *Ap, At*, and *Ai* co-cultures had the largest lag decreases. The *Aa* and *Ao* co-cultures had a smaller, although still significant, decrease (Fig. 4a,b); interestingly, *Aa* and *Ao* were also less efficient at consuming lactate than the other *Acetobacter* species (Fig. 2c). These data indicate that interspecies interactions can change the physiology of the community, and that differences across the acetobacters constitute an opportunity to probe the underlying cause of the lag phenotype.

**Figure 4:**
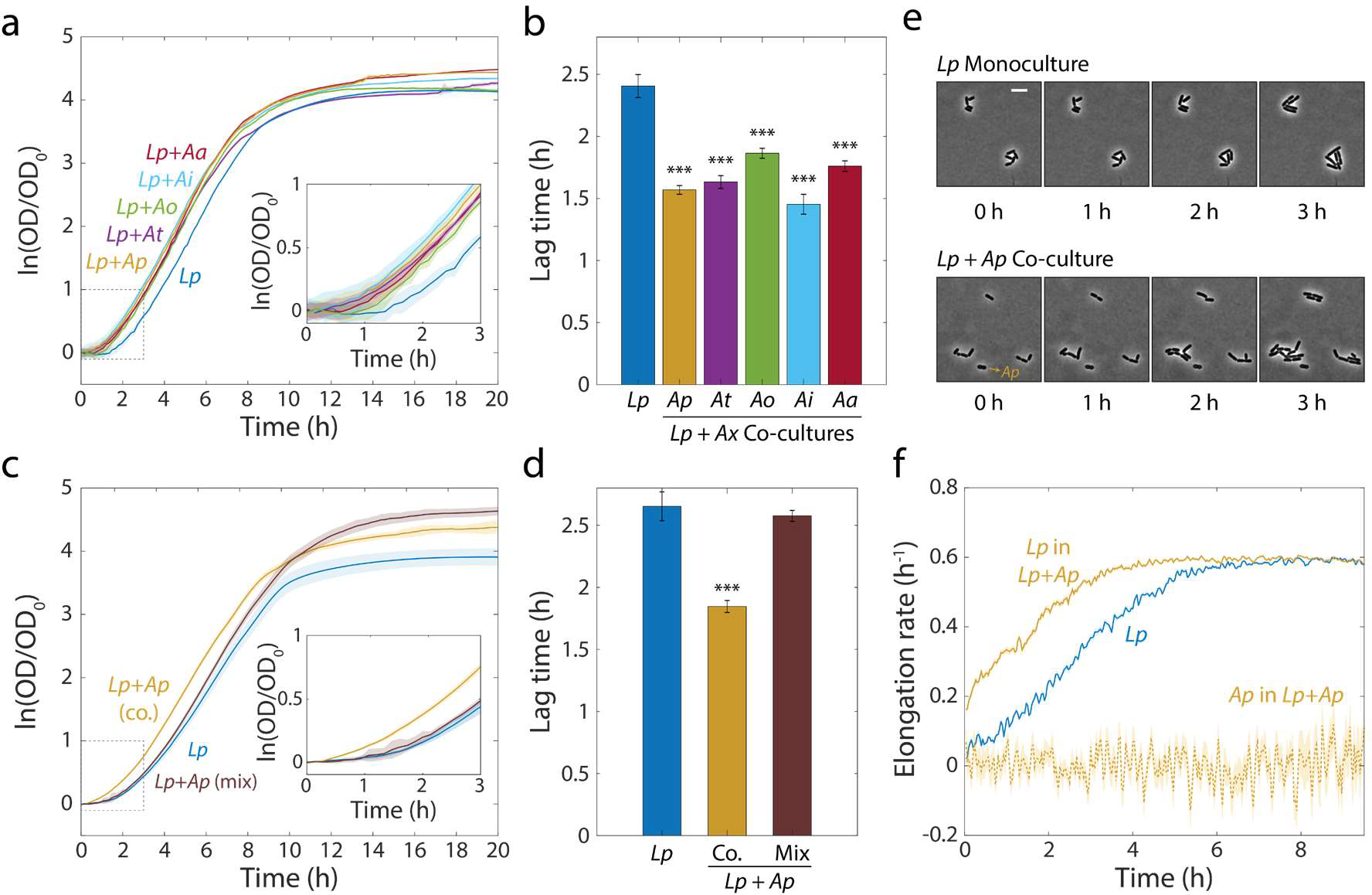
Co-cultures of *Lp* and acetobacters undergo shorter lag phases. a) Calculating the logarithm of OD normalized by OD at *t*=0 reveals that co-cultures of *Lp* and various acetobacters (*A<u>x</u>*) experience more rapid transitions from stationary phase to exponential growth than monocultures of *Lp*. Shaded regions indicate standard deviation (S.D.), *n*=5. Inset: zoom-in of region inside dashed box highlighting lag differences. b) Co-culture lag times are significantly shorter than *Lp* monoculture lag times. Lag times were obtained by fitting the growth curves in (a) to the Gompertz equation. Error bars are S.D. for each condition, *n*=5. *P*-values are from a Student’s two-sided *t*-test of the difference from the monoculture (***: *P*<2×10^−4^). c) Mixing *Lp* monocultures with *Ap* monocultures (Mix) yields growth curves with a similar lag phase than those of *Lp* monocultures. Shaded regions indicate S.D., *n*=5. Inset: zoom-in on region inside dashed box highlighting lag differences. d) Mixed *Lp-Ap* cultures do not experience significantly shorter lag times than *Lp* monocultures. Lag times were obtained by fitting the curves in (c) to the Gompertz equation. Error bars are S.D. for each condition, *n*=5. *P*-values are from a Student’s two-sided *t*-test of the difference from the monoculture (***: *P*<5×10^−4^). e) Single-cell microscopy demonstrates that a decrease in the duration of lag phase of *Lp* was responsible for the lag-time decrease in co-culture. Representative phase microscopy images of *Lp* in monoculture and co-cultured with *Ap* on an MRS agar pad. The only *Ap* cell visible in these images is indicated with an arrow. Size bar = 5 μm. f) The instantaneous elongation rate of single *Lp* cells increases faster in co-culture than in monoculture. Phase-contrast images were segmented and cells were classified as *Lp* or *Ap* based on their aspect ratio. Lines are the mean and shaded regions are the standard error for an *Lp* monoculture (*n*_*Lp,0 h*_ = 465, *n*_*Lp,9.5 h*_ = 27,503) or a co-culture with *Ap* (*n*_*Lp,0 h*_ = 448, *n*_*Lp,9.5 h*_ = 58,087, *n*_*Ap,0 h*_ = 47, *n*_*Ap,9.5 h*_ = 146).

As with *Lp* tolerance to antibiotics (Fig. 1b), the shortened lag phase of the *Lp-Ap* co-culture was history dependent. When we mixed 48-h cultures of *Lp* and *Ap* to a combined initial OD of 0.0375 in the absence of antibiotics, the resulting bulk culture had the same lag time as an *Lp* monoculture (Fig. 4c,d). To determine which of the two species was responsible for the decrease in lag, we diluted a 48-h co-culture of *Lp* and *Ap* 1:200, spotted 2 μL onto a 1% agarose + MRS pad, and performed time-lapse microscopy (Methods) to monitor the initiation of growth at the single-cell level (Fig. 4e). *Lp* and *Ap* are clearly distinguishable based on morphology (Fig. S9): *Lp* cells are longer (2.46 ± 0.78 μm vs. 1.66±0.38 μm) and thinner (0.72 ± 0.12 μm vs. 0.90 ± 0.09 μm) than *Ap* cells. Therefore, we used the aspect ratio (length/width; 3.41 ± 0.91 for *Lp* and 1.87 ± 0.46 for *Ap*) to distinguish single cells from each species in co-culture. We validated this strategy on co-cultures of fluorescently tagged strains of the same two species and observed a 10% error rate in classification (Fig. S9d-f). In co-culture, most *Lp* cells were observed to have grown by 1 h after spotting, but in *Lp* monoculture, few cells were growing even after 2 h (Fig. 4e,f). *Ap* cells did not grow during the time of imaging (Fig. 4f), indicating that the reduced lag time was due to *Lp*’s growth, in agreement with CFU/mL measurements (Fig. 2b).

### Growth status and pH are drivers of antibiotic tolerance

Since pH changes shortened lag phase (Fig. 3e), and since changes in lag time were related to antibiotic tolerance (Fig. 1,4), we tested whether shortening lag phase was sufficient to induce tolerance. Lag phase can be manipulated by increasing time in starvation^21^. To determine the relationship between time spent in stationary phase and lag time in *Lp*, we diluted a 48-h monoculture to a starting OD=0.02 in fresh medium and grew it for varying amounts of time. We then diluted these monocultures into fresh medium at OD=0.0375 and measured bulk culture growth in a plate reader. Incubating the *Lp* monocultures for more than 48 h resulted in a dramatic increase in the duration of lag phase, while reducing the culturing time shortened lag phase (Fig. 5a).

**Figure 5:**
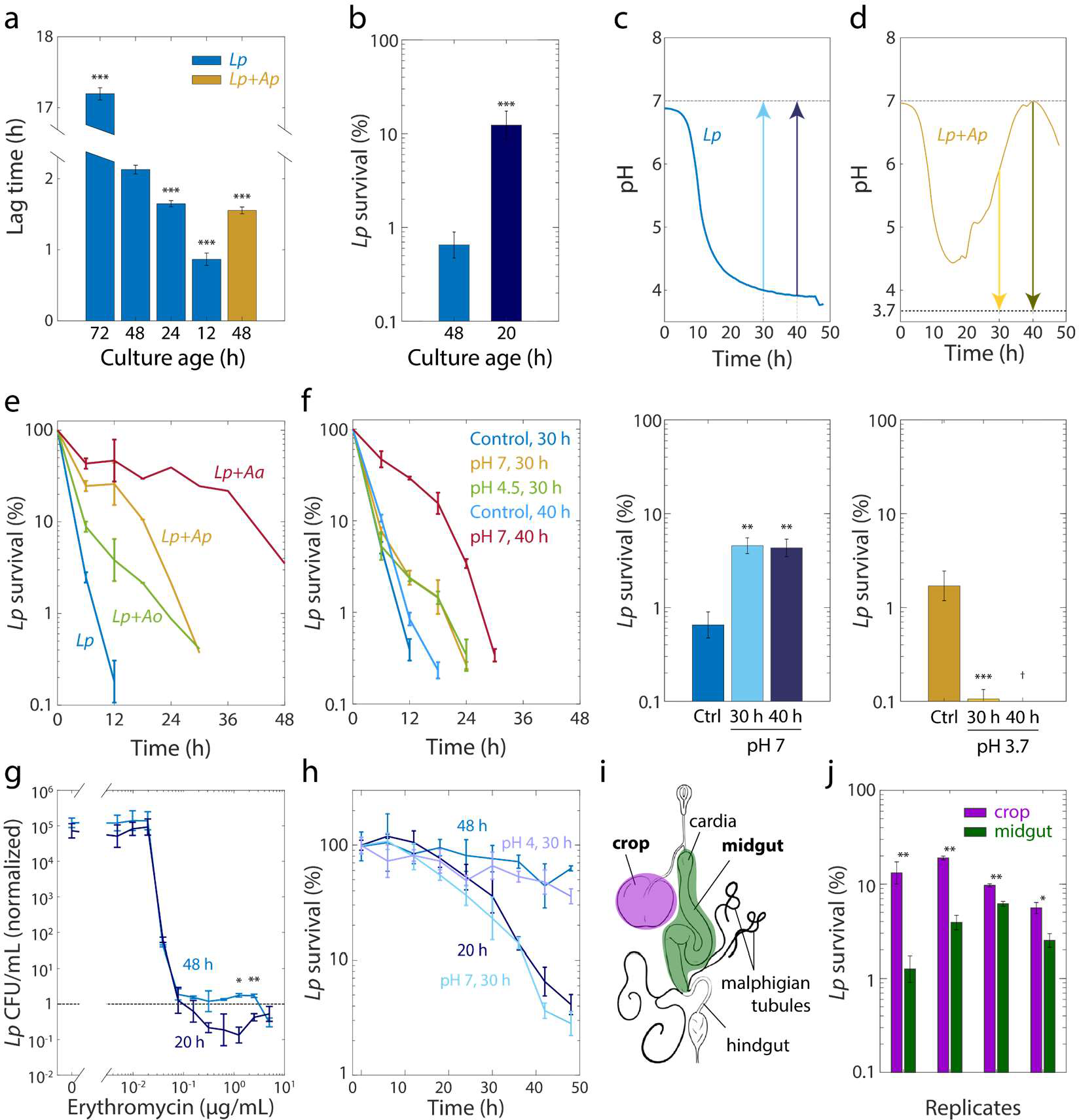
Tolerance to rifampin is modulated by pH. a) The duration of lag phase of bulk cultures of *Lp* depends on the time spent in stationary phase. *Lp* monocultures grown for various times from OD=0.02, and co-cultures with *Ap*, were diluted into fresh medium. Lag time was calculated by fitting growth curves to the Gompertz equation. Error bars are standard deviation (S.D.), *n*=12. *P*-values are from a Student’s two-sided *t*-test of the difference with respect to the 48 h culture (\*\*\**P*<2.5×10^4^). b) Culturing *Lp* as a monoculture for a shorter time leads to higher cell survival. Viable cell plating counts of *Lp* after growth in 20 μg/mL rifampin for 24 h normalized to the counts at the start of the experiment (t=0). Error bars are S.D. for each condition, *n*=3. *P*-values are from a Student’s two-sided *t*-test of the difference between the cultures (***: *P*<1×10^−3^). c) Neutralization of pH in stationary phase in *Lp* monocultures is sufficient to induce tolerance. Increasing the pH of an *Lp* monoculture at t = 30 h or t = 40 h to 7 to mimic the pH increase in co-cultures of *Lp* with acetobacters (upper panel) increased cell survival after treatment with 20 μg/mL rifampin for 24 h (lower panel). A 48-h-old culture with no changes in pH was used as a control (Ctrl.). Error bars are S.D. for each condition, *n*=3. *P*-values are from a Student’s two-sided *t*-test of the difference between the cultures (**: *P*<5×10^−3^). d) Acidification of *Lp* co-cultures with *Ap* during the exponential-to-stationary phase transition or in late stationary phase sensitizes *Lp* to rifampin. Decreasing the pH of an *Lp* co-culture with *Ap* at t = 30 h or *t* = 40 h to 3.7 to mimic the pH of an *Lp* monoculture (upper panel) increased survival after treatment with 20 μg/mL rifampin for 24 h (lower panel). Error bars are S.D. for each condition, *n*=3. *P*-values are from a Student’s two-sided t-test of the difference between the cultures (***: *P*<5×10^−4^). ^†^Values below the limit of detection. e) The dynamics of killing in *Lp* co-culture with acetobacters differs quantitatively according to species and from *Lp* monoculture (blue), indicating that the acetobacters induce rifampin tolerance to different degrees. Normalized CFU/mL of *Lp* in monoculture and in co-culture with acetobacters, and treated with 50 μg/mL rifampin. Error bars are S.D. for each condition, *n*=3. f) The timing of the pH change in Acetobacter co-culture predicts the extent of protection against 50 μg/mL rifampin. Neutralization of pH in *Lp* monocultures at 40 h of growth (to mimic *Lp+Aa* co-cultures) elicits longer protection against rifampin than neutralization at 30 h. A small increase in pH (from 3.85 to 4.5) at 30 h (to mimic *Lp*+Ao co-cultures) provides protection comparable to complete neutralization. Error bars are S.D. for each condition, *n*=3. g) *Lp* survival to erythromycin is ~10 times higher after 24 h of treatment with erythromycin at supraMIC concentrations on 48-h-old monocultures (48 h) of *Lp* than on 20-h-old *Lp* monocultures (20 h). Viable cell plating counts of *Lp* after growth in erythromycin for 24 h normalized to cell counts at the start of the experiment (t=0). Error bars are S.D., n = 3. *P*-values are from a Student’s two-sided *t*-test of the difference between the two samples at a given time point (*: *P*<4×10^−3^, **: *P*<8×10^−4^). h) Shifting the pH of an *Lp* monoculture at 30 h to 4 or 7, followed by 18 h of growth before treatment with 2 μg/mL erythromycin, mimics the survival dynamics of a 48-h-old or 20-h-old culture in stationary phase, respectively. Normalized CFU/mL of *Lp* monocultures. Error bars are S.D. for each condition, *n*=3. i) Schematic of the fruit fly intestinal tract, a ~5 mm-long tube consisting of the foregut, midgut, and hindgut. The crop is an accessory fermentative organ within the foregut. j) Survival of *Lp* is significantly higher in bulk cultures resuspended from the crop than in cultures resuspended from the midgut. Female, ~7-day-old, germ-free flies were colonized with *Lp* and left for three days in sterile food to reach equilibrium, before the crop was dissected from the midgut (Methods). After homogenization of pools of crops and midguts, the cultures were exposed to 50 μg/mL rifampin for 24 h and viable cells were counted via CFU. Error bars are S.D. of the technical replicates for each biological replicate, *n*=3 in each biological replicate. *P*-values are from a Student’s two-sided t-test of the difference between the two samples (*:*P*<1.25×10^−2^, **: *P*<2.5×10^−3^).

Because lag phase in co-culture is slightly shorter than that of a 24-h monoculture (Fig. 5a), we decided to match the lag time of a co-culture by growing a monoculture for 20 h from an initial OD=0.02. We then measured *Lp* survival in 20 μg/mL rifampin after 24 h in cultures diluted from a 48-h-old or a 20-h-old culture. Culturing for 20 h resulted in a significant increase in survival (Fig. 5b). We next tested whether shortening lag phase by changing the pH in stationary phase also yielded increased rifampin tolerance of *Lp* in co-culture. We increased the pH of an *Lp* monoculture to 7 after 30 and 40 h, and grew cells for an additional 18 h and 8 h, respectively. We then measured the change in CFU/mL upon treatment with 50 μg/mL rifampin for 24 h. The upshift in pH at *t* = 30 h or 40 h resulted in increased tolerance relative to the unshifted monoculture (Fig. 5c). To test the extent to which changes in pH affect tolerance, we grew co-cultures of *Lp* and *Ap* for a total of 48 h and decreased the pH to 3.7 at *t* = 30 h or 40 h. In both cases, the viability after 24 h of rifampin exposure was significantly reduced relative to an untreated monoculture (Fig. 5d). Thus, pH can affect tolerance both positively and negatively.

In co-culture with *Lp, Ap* raised the pH earlier than did *Aa*, while *Ao* only raised the pH very slightly (Fig. 2d). We hypothesized that due to these distinct pH dynamics, rifampin would also have different killing *Lp* dynamics in these co-cultures. We grew co-cultures of these acetobacters with *Lp* as previously, and then treated the co-cultures with 50 μg/mL rifampin. While all co-cultures had extended survival relative to *Lp* monoculture, the killing dynamics of *Lp* were indeed distinct, with *Aa* inducing the highest tolerance (Fig. 5e). To determine the extent to which these dynamics can be explained by the time at which each species raises the pH, we measured CFU/mL at various time points after rifampin treatment for *Lp* monocultures grown for 48 h whose pH was raised to pH 7 at *t* = 30 h or 40 h, mimicking the early and late increases in pH for *Ap* and Aa co-cultures, respectively. The shift to pH 7 at 40 h induced higher rifampin tolerance than the shift at 30 h (Fig. 5f), consistent with the increased tolerance of the Lp-Aa co-culture (Fig. 5e). Moreover, shifting the pH to 4.5 at *t* = 30 h, to mimic the slight increase caused by Ao, was also sufficient to increase tolerance comparable to pH neutralization at *t* = 30 h (Fig. 5f), consistent with the similar killing dynamics of the *Ap* and Ao co-cultures (Fig. 5e). All pH shifts induced higher tolerance compared to control cultures that underwent the same protocol but whose pH was maintained (Fig. 5f). Taken together, these experiments establish that pH changes drive the tolerance of *Lp* to rifampin through changes in the exit from stationary phase.

### Growth status and pH also drive tolerance to a ribosome-targeting antibiotic

The robust relationships among changes in pH, lag time, and rifampin tolerance prompted us to explore how changes in pH and lag time affect survival to other antibiotics. We decided to use the ribosome-targeting macrolide erythromycin because it is bactericidal and *Lp* is sensitive to it (Table S2). We treated *Lp* monocultures grown for 20 h or 48 h with increasing concentrations of erythromycin for 24 h at a starting cell density of ~5×10^5^ CFU/mL. In contrast to our observations with rifampin (Fig. 5b), a 48-h *Lp* monoculture displayed tolerance to erythromycin, while a 20-h culture did not (Fig. 5g). While both cultures had the same MIC in erythromycin (0.078 μg/mL, Fig. 5g), at concentrations above the MIC, the 48-h culture showed no changes in CFU/mL after 24 h of erythromycin treatment; the 20-h culture had a reduction of ~10-fold in CFU/mL (Fig. 5g). This result suggests that *Lp* cells diluted from a 48-h culture are tolerant to erythromycin, opposite to the effect we observed with rifampin.

To further determine whether antibiotic tolerance underlies the survival of *Lp* to erythromycin as well as to rifampin, we diluted 20- and 48-h cultures to a starting density of ~5×10^5^ CFU/mL, exposed them to a high concentration of erythromycin (2 μg/mL, 25X MIC), and monitored CFU/mL over time. Cells from a 20-h culture died significantly more rapidly than cells from a 48-h-old culture (Fig. 5h), indicating that the differences in survival (Fig. 5g) are explained by erythromycin tolerance. Further, increasing the pH of an *Lp* monoculture at 30 h and then exposing it after an additional 18 h of growth to 2 μg/mL erythromycin in fresh medium had an increase in the rate of killing that was similar to that achieved with a 20-h culture (Fig. 5h). These results highlight that the effects of changing the growth status of a culture by different means are not limited to rifampin and—as in the case of erythromycin—can be opposite.

### *Growth state in the fruit fly gut determines antibiotic tolerance* ex vivo

Our *in vitro* observations connecting changes in pH to lag time and antibiotic tolerance prompted us to examine whether these properties are also linked within the fruit fly gut. This tract consists of a ~5-mm-long tube divided into three sections: foregut, midgut, and hindgut (Fig. 5i). The foregut includes an accessory storage organ known as the crop. The contents of the crop are delivered to the midgut through the proventriculus and transit through the midgut to end in the hindgut, where they are expelled into the environment through the rectum and the anus^22^. Specialized “copper” cells in the central portion of the midgut keep the pH of this section low, akin to the stomach in mammals^23^. Bacteria are distributed along the *Drosophila* gastrointestinal tract; *Lp* in particular can colonize all compartments with slight variations in its distribution along the tract^24^.

To examine whether variations along the fly gut impact *Lp* physiology, we coarse-grained the digestive tract into the crop and the midgut and quantified rifampin tolerance of *Lp* from these regions. We hypothesized that the different functions of these regions - storage, or nutrient absorption and transit - lead to differences in bacterial physiology. We colonized 5-7-day-old, female, germ-free flies with *Lp* and left them for three days in sterile food to reach equilibrium (Methods). We then dissected the flies and separated the crop from the midgut. We pooled (i) dissected crops and (ii) dissected midguts in MRS to obtain a final bacterial density of ~5×10^5^ CFU/mL. Crops had overall ~10 times less *Lp* than the midgut (8,700 CFU/crop vs. 86,000 CFU/midgut). After homogenization, the samples were exposed to 20 μg/mL or 50 μg/mL rifampin for 24 h. The MIC of rifampin for these *ex vivo* samples was approximately the same as for *in vitro* cultures (1.25 μg/mL). For cultures extracted from the crop, significantly more cells survived than for cultures extracted from the midgut, with values comparable to those of a co-culture with *Ap* and a monoculture, respectively (Fig. 5j). These results indicate that spatial heterogeneity in the host can lead to differences in the duration of lag phase and antibiotic tolerance *ex vivo*.

## Discussion

Our measurements of the growth behavior of the fly gut microbiota indicate that interspecies interactions impact both the metabolism of a microbial community and the effect of antibiotics on individual species. For fly gut commensals, the pH-based mechanism underlying the tolerance of *Lp* induced by acetobacters is intrinsically connected to the metabolic capacity of each species, and hence is likely to be generally relevant *in vivo* to the resilience of this community under perturbations. Moreover, these findings could have important implications for human health, for example in the context of *Lactobacillus*-dominated vaginal microbiotas^25^, and their generality should be tested broadly in other contexts.

In this study, we observed a novel form of antibiotic tolerance. Tolerance has been defined as increased time to killing^26^, as opposed to resistance (a change in the MIC), or persistence (the ability of a subpopulation of clonal bacteria to survive high concentrations of antibiotic^5^). Tolerance to beta lactams such as ampicillin has been observed in *E. coli* cultures that exhibit slow growth or a long lag phase^5^, and *E. coli* mutants with longer lag phases can be selected through experimental evolution to match the time of treatment^27,28^. Based on these previous studies, we were surprised to find the opposite effect with rifampin on *Lp*: cultures with a shorter lag phase exhibited increased tolerance (Fig. 1,4).

Moreover, although tolerance to erythromycin was associated with a longer lag phase (Fig. 3e, 5a,h), killing retardation was at least an order of magnitude longer than the change in lag time (Fig. 5a,h), indicating that tolerance is not determined by an elongation of lag phase alone, in contrast to the effects of ampicillin on *E. coli*^27^.

Several genetic factors that increase time to killing have been identified in *E. coli*, including toxin-antitoxin modules such as *hipBA*^29^ that induce the stringent response and thus cause transient growth arrest. In *Lp* co-culture with acetobacters, metabolic interactions alter the physiological state of *Lp* during late stationary phase by changing the environmental pH (Fig. 2). The stringent response is required to survive acid shock in *Helicobacter pylori*^30^ but not in *Enterococcus faecalis*^31^, which is in the same order as *Lp*. In the case of *Lp*, whether the stringent response could be a major factor in the increased tolerance to rifampin is unclear due to the surprising connection with decreased lag (not to mention the opposite behavior with erythromycin).

The pH in stationary phase can affect many factors, such as the chemistry of extracellular metabolites and macromolecules as well as the surface of the cell^32^. Importantly, our assays of antibiotic sensitivities were all performed at a starting pH of 7. Nonetheless, shifts in extracellular pH can lead to buffered drops in cytoplasmic pH^33,34^; such drops can be regulated^35^ or result from internalization of low-p*K*_a_ species such as short-chain fatty acids^36^. Such changes could lead to protonation of macromolecules involved in adsorption or changes in the proton motive force^37^. How these factors affect non-polycationic antibiotics such as rifampin remains to be determined; neither of the ionizable functional groups of rifampin (p*K*_a_s 1.7 and 7.9^38^) nor erythromycin (p*K*_a_ 8.88^39^) have p*K*_a_s in the pH range achieved in our cultures (Fig. 2,S7b). Protonation changes in target macromolecules could also lead to protection against antibiotics, although we would expect a subsequent change in MIC, contrary to our findings (Fig. 1,5). Intracellular acidification by the short-chain fatty acid propionate has been proposed to lengthen lag phase in *Salmonella in vitro* and in the mouse gut^40^, consistent with our finding that lag time (Fig. 4) and intracellular pHluorin fluorescence (Fig. S6a) are related.

Changes in intra- and extracellular pH have been shown to lead to transcriptional responses that provide cross-protection against antibiotics^41–43^, suggesting that the killing retardation due to a pH increase in stationary phase may result from a complex regulatory process. One major factor influencing the *Lactobacillus-Acetobacter* interaction is that these organisms form a recurrent community and may therefore have evolved to sense and benefit from each other’s presence. Further experiments are needed to uncover the molecular mechanisms that link growth state and susceptibility to antibiotics in lactobacilli, other non-model organisms, and microbial communities. In addition, although we consistently observed related shifts in lag phase and tolerance (Fig. 3,5), it remains to be established whether lag time and tolerance are causally linked or coupled to some global variable, particularly given the opposite effects on rifampin and erythromycin tolerance.

Previous work has shown that bacterial interactions can elicit changes in antibiotic sensitivity by changing cellular physiology or interfering with antibiotic action directly or indirectly^9–11^. In principle, a myriad of intra- and extra-cellular variables are subject to the composition and dynamics of the ecosystems that bacteria inhabit, and microbial communities within mammalian hosts can elicit changes in environmental variables both locally and globally. Specifically, the microaerobic and anaerobic microenvironments of the human and fly^24^ gastrointestinal tracts enable the growth of short chain fatty acid producers. Some of these short chain fatty acids, like butyrate, have been shown to play an important role on host physiology and health^44^. The consequences of the accumulation of these short chain fatty acids and other small molecules on microenvironments, as well as their effect on bacterial physiology and antibiotic treatment efficacy *in vivo*, have yet to be systematically explored. Our results emphasize the need to probe the action of antibiotics – as well as other drugs that are thought not to target microbial growth^45^ – in complex and varied conditions^46^. Furthermore, our findings highlight the utility of studying growth physiology in co-cultures in the absence of antibiotics for uncovering novel mechanisms of community-encoded protection against antibiotics.

## Online Methods

### Fruit fly stocks and gut microbiome sequencing

*Wolbachia*-free *Drosophila melanogaster* Canton-S (BL64349) flies were obtained from the Bloomington Drosophila Stock Center, and were reared and maintained as previously described^24^. To determine the bacterial strains present in our flies, we performed culture-independent 16S amplicon sequencing targeting the V4 region on an Illumina MiSeq. Individual flies were CO_2_-anesthetized, surface-sterilized by washing with 70% ethanol and sterile PBS six times each. Flies were dissected under a stereo microscope and their guts were placed in 2-mL screw cap microtubes containing 200 μL of 0.1-mm sterile zirconia-silicate beads (BioSpec Products 11079101z) and 350 μL of sterile lysis buffer (10 mM Tris-HCl, pH 8, 25 mM NaCl, 1 mM EDTA, 20 mg/mL lysozyme). Samples were homogenized by bead beating at maximum speed (Mini-Beadbeater, BioSpec Products) for 1 min. Proteinase K was added at 400 μg/mL and samples were incubated for 1 h at 37 °C. Sample were then centrifuged (3,000 × *g* for 3 min) and 300 *μ*L of the nucleic acids-containing supernatant were transferred to 1.7-mL microtubes. Genomic DNA from samples was cleaned up through a DNA Clean & Concentrator-5 column (Zymo Research D4014). Using the protocol described in Ref. ^47^ for library preparation and sequencing, we sequenced the gut contents of 18 individual flies, three flies each from six independent vials. Paired-end 250-base pair sequencing generated >10,000 reads per sample. Reads were filtered using PrinSeq as in Ref ^48^. The reads were then clustered into operational taxonomic units (OTUs) at 99% identity and assigned taxonomy using LOTUS^49^ with the following parameters: [-threads 60 -refDB SLV -highmem 1 -id 0.99 -p miseq-useBestBlastHitOnly 1 -derepMin 3:10,10:3 -simBasedTaxo 1 -CL 3]. Redundant strain identities were collapsed into single OTUs. Common reagent contaminant strains were then removed^50^. After filtering, only five unique species were identified (Fig. 1a). We isolated these species in culture and verified the taxonomic identity of our isolates using Sanger sequencing of the complete 16S rRNA gene^13^. At 97% OTU clustering, only three species were found: *Acetobacter* sp., *Lactobacillus plantarum*, and *Lactobacillus brevis*. When less stringent FASTQ quality filtering was used, trace amounts (~0.01%) of two mammalian gut strains were identified: *Blautia* sp. and *Bacteroides* sp. Because these OTUs were eliminated by more stringent quality filtering, we speculate that they may have resulted from barcode bleed-through on the MiSeq flowcell.

### Bacterial growth and media

Bacterial strains used in this study are listed in Supplementary Table 1. For culturing, all strains were grown in MRS medium (Difco™ Lactobacilli MRS Broth, BD 288110). Frozen stocks were streaked onto MRS agar plates (1.5% agar, Difco™ agar, granulated, BD 214530) and single colonies were picked to start cultures. MYPL medium was adapted from Ref. ^51^, with 1% (w/v) D-mannitol (ACROS Organics AC125345000, Lot A0292699), 1% (w/v) yeast extract (Research Products International Y20020, Lot 30553), 0.5% (w/v) peptone (Bacto™ peptone, BD 211677 Lot 7065816), 1% (w/v) lactate (Lactic acid, Sigma L6661-100ML Lot MKCC6092), and 0.1% (v/v) Tween^®^ 80 (Polyoxyethylene(20)sorbitan monooleate, ACROS Organics AC278632500 Lot A0375189). The medium was set to pH 7 with NaOH (EMD Millipore SX0590, Lot B0484969043). All media were filter-sterilized. Strains were grown at 30 °C with constant shaking.

To count CFUs in cultures, aliquots were diluted serially in PBS. For cultures treated with high concentrations of antibiotics, cells were centrifuged for 1.5 min at 8000 x *g* and resuspended in 1X PBS pH 7.4 (Gibco™ 70011044) after removing the supernatants. PBS-diluted cultures were plated on MRS and MYPL because lactobacilli grow faster than acetobacters on MRS and vice versa on MYPL. Colony morphology and color enable differentiation of lactobacilli from acetobacters.

### Conditioned media

Conditioned media were obtained by centrifuging cultures at 4500 x *g* for 5 min and filtering the supernatant with a 0.22-μm polyethersulfone filter (Millex-GP SLGP033RS) to remove cells. Conditioned media were acidified with HCl (Fisher Chemical A144-500, Lot 166315) or basified with NaOH (EMD Millipore SX0590, Lot B0484969043). Conditioned media were sterilized after adjusting pH with 0.22-μm PES filters.

### MIC estimations

To estimate the sensitivity of each species to various antibiotics, colonies were inoculated into MRS and grown for 48 h at 30 °C with constant shaking. Cultures were diluted to an OD of 0.001 for *Lp*, *Lb*, and *At*, and 0.01 for *Ap*. Diluted cultures (195 μL) were transferred to 96-well plates containing 5 μL of antibiotics at 40X the indicated concentration. Antibiotics used were ampicillin (ampicillin sodium salt, MP Biomedicals 02194526, Lot R25707, stock at 100 mg/mL in milliQ H_2_O), streptomycin (streptomycin sulfate salt, Sigma S9137 Lot SLBN3225V, stock at 50 mg/mL in milliQ H_2_O), chloramphenicol (Calbiochem 220551, Lot D00083225, stock at 50 mg/mL in ethanol), tetracycline (tetracycline hydrochloride, MP Biomedicals 02103011, Lot 2297K, stock at 25 mg/mL in dimethyl sulfoxide (DMSO)), erythromycin (Sigma E5389-1G, Lot WXBC4044V, stock at 64 mg/mL in methanol), ciprofloxacin (Sigma-Aldrich 17850, Lot 116M4062CV, stock at 1.2 mg/mL in DMSO), trimethoprim (Alfa Aesar J63053-03, Lot T16A009, stock at 2 mg/mL in DMSO), spectinomycin (spectinomycin hydrochloride, Sigma-Aldrich PHR1426-500MG, Lot LRAA9208, stock at 50 mg/mL in milliQ H_2_O), rifampin (Sigma R3501-5G, Lot SLBP9440V, stock at 50 mg/mL in DMSO), and vancomycin (vancomycin hydrochloride, Sigma-Aldrich PHR1732-4X250MG, Lot LRAB3620, stock at 200 mg/mL in DMSO:H_2_O 1:1). Antibiotics were diluted serially in 2-fold increments into MRS. Cultures were grown for 24 h at 30 °C with constant shaking and absorbance was measured in an Epoch2 plate reader (BioTek Instruments) at 600 nm. The MIC was estimated as the minimum concentration of antibiotic with absorbance within two standard deviations of media controls.

For experiments in Supplementary Figure S2, colonies of *Lp* and *Ap* were inoculated into MRS and grown for 48 h at 30 °C with constant shaking. The saturated cultures were diluted to OD 0.02, mixed 1:1, and grown for 48 h at 30 °C with constant shaking. Then, the mono- and co-cultures were diluted to an OD of 0.001 (final cell density ~5×10^5^ CFU/mL) and transferred to 96-well plates containing 5 μL of rifampin at 40X working concentration. Cultures were grown for 24 h and MICs were estimated as described above. For Figures 1b-e, cultures were serially diluted in 5-fold increments in PBS, and 3 μL of the dilutions were spotted onto MRS and MYPL rectangular plates using a semi-automated high-throughput pipetting system (BenchSmart 96, Mettler Toledo). Plates were incubated at 30 °C until colonies were visible for quantification of viability.

### Plate reader growth curves

Cultures were grown from single colonies for 48 h in MRS at 30 °C with constant shaking. Then, cultures were diluted to a final OD of 0.02 and 200 μL of the dilutions were transferred to clear-bottom transparent 96-well plates. Plates were sealed with transparent film pierced with a laser cutter to have ~0.5-mm holes to allow aeration in each well. Absorbance was measured at 600 nm in an Epoch2 plate reader (BioTek Instruments). Plates were shaken between readings with linear and orbital modes for 145 s each.

Growth rates and lag times were quantified using custom MATLAB (Mathworks, R2008a) code. The natural logarithm of OD was smoothed with a mean filter with window size of 5 timepoints for each condition over time, and the smoothed data were used to calculate the instantaneous growth rate *d*(ln(OD))/*dt*. The smoothed ln(OD) curve was fit to the Gompertz equation^52^ to determine lag time and maximum growth rate.

### pH measurements

Culture pH was measured using the dual-excitation ratiometric pH indicator 2’,7-bis-(2-carboxyethyl)-5-(and-6)-carboxyfluorescein, mixed isomers (BCECF, Invitrogen B1151, Lot 1831845), which has a p*K*_a_ of ~6.98. A stock solution of 1 mg/mL BCECF in DMSO (Fisher BioReagents BP231, Lot 165487) was diluted 1000-fold into MRS to a final concentration of 1 μg/mL. Cells were grown in a Synergy H1 plate reader (BioTek Instruments) following the procedure described above. In addition to absorbance, fluorescence was measured every cycle using monochromators at excitation (nm)/emission (nm) wavelength combinations 440/535 and 490/535. After subtracting the fluorescence of wells containing cells without the indicator, the ratio of the signals excited at 440 nm and 490 nm was used to calculate the culture pH using a calibration curve of MRS set to various pH values.

Culture pH after 48 h of growth was directly measured with a pH meter (sympHony, VWR) equipped with a pH combination electrode (Fisherbrand™ accumet™ 13-610-104A).

### Changes in pH during growth

To change the pH of monocultures and co-cultures in stationary phase, we obtained conditioned medium at 30 h or 40 h as described above and set the pH to the desired values. We then centrifuged 2 mL of a replicate culture for 3 min at 8000 x *g*, removed the supernatant, and resuspended cells in 1 mL of the corresponding medium to wash the cells. The suspension was centrifuged a second time and the pellets were resuspended in 2 mL of the corresponding medium.

### Time-lapse and fluorescence microscopy

Cells were imaged on a Nikon Eclipse Ti-E inverted fluorescence microscope with a 100× (NA 1.40) oil-immersion objective. Images were collected on a DU897 electron multiplying charged couple device camera (Andor) using μManager v.1.4^53^. Cells were maintained at 30 °C during imaging with an active-control environmental chamber (Haison).

Cultures grown for 48 h were diluted 100-fold into PBS and 2 μL were spotted onto a 1% (w/v) agarose MRS pad. After drying at room temperature, the pads were covered with a cover slip, sealed with a mixture of equal portions of Vaseline, lanolin, and paraffin, and transferred to the microscope. Images were taken every 2 min using μManager v. 1.4.

To quantify the morphology of cells using fluorescent strains, co-cultures were diluted 100-fold into PBS and 2 μL were spotted onto a 1% (w/v) agarose PBS pad. After drying, the pads were covered with a cover slip and transferred to the microscope. Images were acquired at room temperature using μManager v. 1.4.

For Figure S6, saturated *Ap* monocultures were diluted 100- or 500-fold into PBS and 2 μL were spotted onto a 1% (w/v) agarose MRS pad containing 30 μM propidium iodide (from a 4.3-mM stock in water, BD, Cell Viability Kit 349483). After drying, the pads were covered with a cover slip and transferred to the microscope. Images were taken at 30 °C every 5 min using μManager v. 1.4.

The MATLAB image processing software *Morphometrics*^54^ was used to segment cells and to identify cell contours from phase-contrast images. Fluorescence intensity per cell was calculated by averaging the fluorescence over the area of the cell. A threshold for propidium-iodide labeling was defined that clearly separated labeled cells from unlabeled cells (data not shown).

### Single-cell tracking and analysis

Images were segmented and cells were tracked using the software *SuperSegger* v. 3^55^. Further analysis of single-cell growth was performed using custom MATLAB code. Cells with length >6 μm were removed from further analysis due to issues with segmentation. Length traces were smoothed using a mean filter of window size 5. Cells were classified as *Lp* or *Ap* if 90% of their traces were above (*Lp*) or below (*Ap*) a log10(length-to-width ratio) of 0.375. Traces with more than 15 timepoints were used for further analysis. Elongation rates *d*(ln *L*)/*dt* were calculated for each cell and the mean and standard error were computed for each time point.

### Cloning and transformations

To generate the fluorescently labeled *Ap* strain, the sfGFP coding sequence was cloned into pCM62^56^ under control of the *Escherichia coli* lac promoter. The sfGFP coding sequence was amplified from pBAD-sfGFP using primers ZTG109 (5’ ggatttatgcATGAGCAAGGGCGAGGAG) and ZTG110 (5’-gctttgttagcagccggatcgggcccggatctcgagTTACTTGTACAGCTCGTCCATG).

Gibson assembly^57^ was used to insert the amplified sfGFP cassette into BglII/XhoI-digested pCM62. This construct was delivered into *Ap* by conjugation as previously described^58^. *Escherichia coli* BW29427 was used as a donor strain and maintained with 80 mg/mL 2,6-Diaminopimelic acid (Sigma Aldrich 33240)in potato agar mating plates^58^. Transformed *Ap* was selected with 10 μg/mL tetracycline on yeast peptone glycerol agar plates^58^.

To generate the *Lp* strain harboring pHluorin, the pHluorin coding sequence was cloned into pCD256-mCherry^59^ under the control of the strong p11 promoter^20^. The pHluorin coding sequence was amplified using primers ZFH064-pHluorin (5’-ATTACAAGGAGATTTTACAT ATGAGTAAAGGAGAAGAACTTTTC) and ZFH065-pHluorin (5’-gtctcggacagcggttttGGATCCTTATTTGTATAGTTCATCCATG). Gibson assembly^57^ was used to insert the amplified pHluorin cassette into NdeI/BamHI-digested pCD256-mCherry. The *Lp*-pHluorin strain was generated by transforming wild type *Lp* as previously described^60^.

Fluorescent strains were further grown in MRS with antibiotics (10 μg/mL chloramphenicol (Calbiochem 220551, Lot D00083225) for *Lp*, tetracycline (10 μg/mL tetracycline hydrochloride, MP Biomedicals 02103011, Lot 2297K) for *Ap*).

### pHluorin measurements

Cells were grown following the procedure described above. The *Lp* pHluorin strain was grown in MRS containing 10 μg/mL chloramphenicol for the first 48 h of growth. In addition to absorbance, fluorescence was measured every cycle using monochromators at excitation (nm)/emission (nm) wavelength combinations 405/509 and 475/509. Because the signal from excitation wavelength 405 nm was undistinguishable from signal from medium (data not shown), we also measured pHluorin signal at both excitation/emission wavelength combinations for cells in PBS. Cultures (48-h-old, 250 μL) were centrifuged at 10,000 x *g* for 1 min and resuspended in 1X PBS. Aliquots (200 μL) were transferred to a 96-well plate and fluorescence was measured using monochromators at excitation (nm)/emission (nm) wavelength combinations 405/509 and 475/509 within 1 min of resuspension in a Synergy H1 plate reader (BioTek Instruments).

### Lactate measurements

Colonies of *Lp* and acetobacters were inoculated into 3 mL MRS and grown for 48 h at 30 °C with constant shaking. Saturated cultures were diluted to OD 0.02, mixed 1:1, and grown at 30 °C with constant shaking. After mixing for 20 h and 48 h, a 700-μL aliquot was transferred to a microcentrifuge tube and centrifuged at 10,000 x *g* for 4 min. Supernatant (600 μL) was transferred to a new tube and centrifuged at 10,000 x *g* for 4 min. Supernatant (500 μL) was transferred to a new tube and kept on ice for not longer than 1 h, until lactate was measured.

L- and D-lactate concentrations were measured using the EnzyChrom™ L-(BioAssay Systems ECLC-100, Lots BH06A30 and BI07A09) and D-lactate (BioAssay Systems EDLC-100, Lots BH0420 and BI09A07) Assay Kits. Samples were diluted 10- and 100-fold in water, and absorbance was measured according to the manufacturer’s instructions in a plate reader (Tecan M200). We also included controls without lactate dehydrogenase to account for endogenous activity in the supernatants.

### Ex vivo *experiments*

We generated germ-free flies by sterilizing dechorionated embryos. Embryos oviposited on grape juice-yeast medium (20% organic grape juice, 10% active dry yeast, 5% glucose, 3% agar) were harvested and washed twice with 0.6% sodium hypochlorite for 2.5 min each, once with 70% ethanol for 30 s, and three times in sterile water for 10 s each. Eggs were transferred into flasks with sterile glucose-yeast medium (10% glucose, 5% active dry yeast, 1.2% agar, 0.42% propionic acid) and were maintained at 25 °C with 60% humidity and 12 h light/dark cycles. Germ-free stocks of these flies were kept for several generations and were regularly checked for sterility by plating flies onto MRS and YPD media.

To prepare flies for tolerance measurements, we took ~3-day-old germ-free flies and transferred them to sterile vials with ~50 flies each. We added ~5 × 10^6^ CFU of *Lp* onto the food and let the flies equilibrate with the bacteria for 3 days. The day before the experiment, flies were transferred into a clean sterile vial.

To extract the midgut and crop, flies were washed with 70% ethanol and PBS six times each. Flies were dissected under a stereo microscope in sterile PBS. Dissected crops and midguts were pooled into 2 mL of sterile MRS with 200 μL 0.5-mm diameter sterile zirconia-silicate beads (BioSpec Products 11079105). Suspended organs were homogenized in a bead beater (Mini-Beadbeater, BioSpec Products) at maximum speed for 1 min. The homogenate was diluted to a cell density of ~5 × 10^5^ CFU/mL and was treated with 50 μg/mL rifampin for 24 h.

### Statistical analyses

To determine significance of differences, we performed pairwise Student’s two-sided *t*-tests throughout. To decrease Type I error, we performed Bonferroni corrections for each experiment. Significant differences are denoted in the figures: *: *P*<0.05/*n*, **: *P*<0.01/*n*, ***: *P*<0.001/*n*, where *n* is the number of comparisons.

## Acknowledgments

The authors thank Vivian Zhang for technical support, Elizabeth Skovran for kindly providing the pCM62 plasmid for *Acetobacter* spp., and Kazunobu Matsushita for providing the *A. tropicalis* SKU1100 control strain in the initial conjugation experiments. We also thank the Huang and Ludington labs for fruitful discussions. This work was supported by NIH Director’s New Innovator Awards DP2OD006466 (to K.C.H.), NSF CAREER Award MCB-1149328 (to K.C.H.), the Allen Center for Systems Modeling of Infection (to K.C.H.), and NIH Director’s Early Independence Award DP5OD017851 (to W.B.L.). K.C.H. is a Chan Zuckerberg Biohub Investigator. A.A.-D. is a Howard Hughes Medical Institute International Student Research fellow and a Stanford Bio-X Bowes fellow.

## Author Contributions

A.A.-D., K.C.H., and W.B.L designed the research. A.A.-D. and T.T. performed *in vitro* experiments. A.A.-D. and B.O. performed *ex vivo* experiments. B.O., Z.T.G., and Z.H. built fluorescent strains. A.A.-D. analyzed the data. A.A.-D., K.C.H., and W.B.L. wrote the paper. All authors reviewed the paper before submission.

