## Supplemental Information for "Bacterial interspecies interactions modulate pH-mediated antibiotic tolerance in a model gut microbiota"

Andrés Aranda-Díaz[1], Benjamin Obadia[2], Tani Thomsen[1], Zachary F. Hallberg[3], Zehra Tüzün Güvener[2], Kerwyn Casey Huang[1,4,5,*], William B. Ludington[2,6,*]

[1]Department of Bioengineering, Stanford University, Stanford, CA 94305, USA

[2]Department of Molecular and Cell Biology, University of California, Berkeley, Berkeley, CA 94720, USA

[3]Department of Plant and Microbial Biology, University of California, Berkeley, Berkeley, CA 94720, USA

[4]Department of Microbiology and Immunology, Stanford University School of Medicine, Stanford, CA 94305, USA

[5]Chan Zuckerberg Biohub, San Francisco, CA 94158

[6]Department of Embryology, Carnegie Institution of Washington, Baltimore, MD 21218

### Supplementary Text

Lp *has a strong positive interaction with* Ap

We grew *Lp* and each of the acetobacters separately for 48 h, diluted the monocultures to OD = 0.04, combined the *Lp* monoculture 1:1 with each *Acetobacter* monoculture, and grew the co-cultures for 48 h in a plate reader. We then diluted the saturated cultures 1:30 in phosphate-buffered saline (PBS) to accurately measure the final OD, and computed an interaction score based on an additive model:

$$\alpha = \frac{\mathrm{OD}_{\mathrm{co}} - (\mathrm{OD}_{Lp} + \mathrm{OD}_{A})}{\sqrt{\mathrm{OD}_{Lp}\mathrm{OD}_{A}}} \quad (1)$$

where $\mathrm{OD}_{\mathrm{co}}$ and $\mathrm{OD}_{A}$ are the final ODs of the co-culture and the *Acetobacter* monoculture, respectively. With this metric,  indicates synergy and  indicates antagonism. *Ap* showed a strong (positive) interaction with *Lp*, whereas the *At*, *Ai*, and *Aa* had an interaction score closer to 0 (Fig. S3a). Only the *Lp*-*Ao* co-culture was significantly antagonistic (Fig. S3a). We also performed this measurement in cultures grown in test tubes, where we only observed a significant positive interaction of *Lp* with *Ap* (Fig. S3a). Calculating the interaction score described above using CFU/mL instead of OD revealed a positive interaction of *Lp* with *Ap* and negative interactions with the rest of the acetobacters (Fig. S3c).

*Cell death of* Ap *during initial outgrowth in MRS depends on the inoculum*

Given the low level of growth of the *Ap* monoculture (Fig. 3b), we tested how growth was affected by the initial inoculum of *Ap*. We grew a 48-h monoculture of *Ap* in MRS, diluted it to various degrees into fresh MRS, and monitored *Ap* growth over time in a plate reader.

Interestingly, the final OD of these cultures was inoculum-dependent (Fig. S6a,b); inocula approximating the $10^5$ CFU/mL values (OD ~ $10^{-3}$) that led to long lag phases and low carrying capacity in our previous experiments (Fig. 3b), had 40% lower carrying capacity than the highest-density inocula (Fig. S6a). Washing the cultures with PBS before diluting them in fresh medium made the inoculum dependence slightly stronger (Fig. S6a,b), although this effect could be due to loss of cells during washing.

To interrogate the cause of this inoculum-dependent growth, we placed 2 μL of high (OD = 0.036, from a saturated culture diluted in PBS) or low inoculum (OD = 0.0072) onto a 1% agarose + MRS pad containing 4.3 mM propidium iodide (PI, a stain generally considered to be taken up only by dead cells) and monitored growth at the single-cell level using time-lapse microscopy. We only observed a few growing cells (<5%) after ~3.5 h of imaging (data not shown). The degree of death was much higher for the low-density inoculum than for the high-density inoculum (30% vs. <1%, Fig. S6c), indicating that the protective effect of inoculum density is due to a reduction in cell death. The growth yield on MRS was not inoculum-dependent for the other acetobacters (Fig. S6d), with the exception of a subtle inoculum-dependency for *Aa* (Fig. S6d), nor was there inoculum dependence for *Ap* on MYPL (Fig. S6a), indicating that this inoculum dependence is not universal or integral to the antibiotic tolerance phenotype.

**Supplementary Figures**

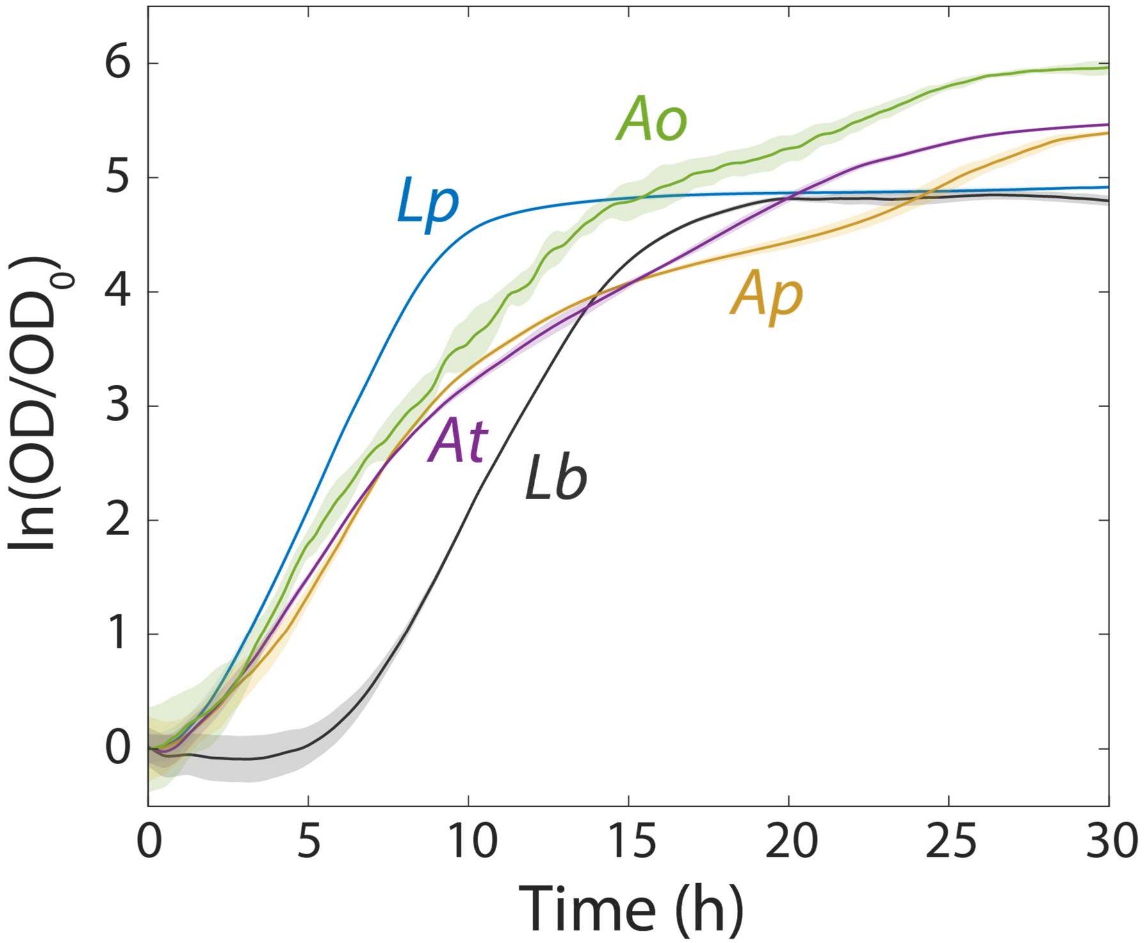


**Supplementary Figure S1: Growth curves of monocultures of the primary species in the fruit fly gut microbiota measured with a plate reader (Methods).** Data were normalized by the initial OD ($t$=0) for each curve. *Lactobacillus plantarum, Lp*; *L. brevis, Lb*; *Acetobacter pasteurianus, Ap*; and *A. tropicalis, At*. Shaded regions are standard deviation, $n$=16.

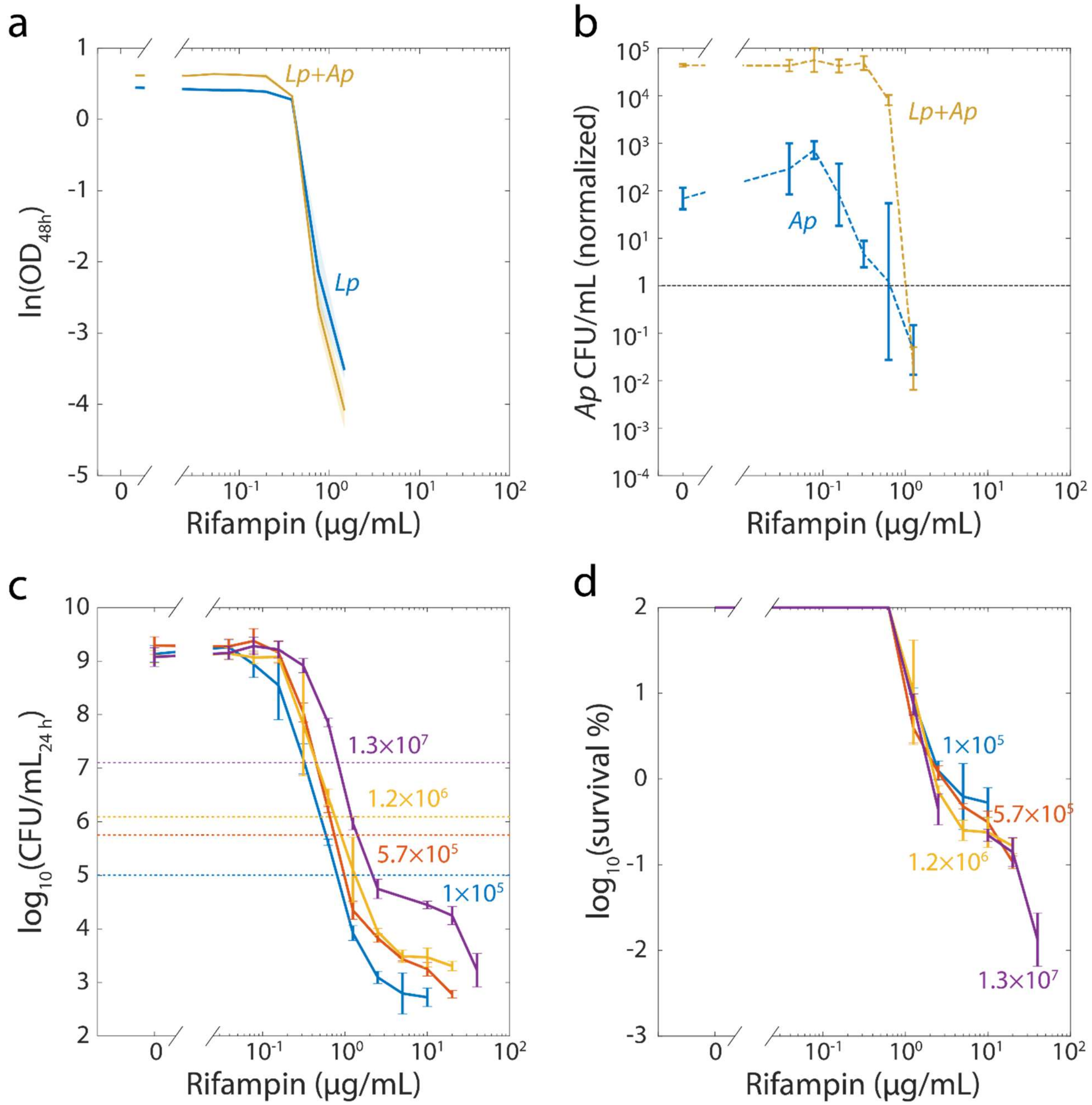


**Supplementary Figure S2: Changes in the survivability of *Lactobacillus plantarum* (*Lp*) when co-cultured with *Acetobacter pasteurianus* (*Ap*) are not due to a change in minimum inhibitory concentration (MIC), increased survivability of *Ap*, or inoculum size.**

a) The MICs of *Lp* in monoculture and in co-culture with *Ap* are the same. Shaded regions are standard deviation (S.D.), $n$=3. The highest concentration of rifampin tested was 40 μg/mL. Data points not shown were below the limit of detection of the plate reader.

b) *Ap* and *Lp* die at similar concentrations of rifampin. Viable cell plating counts of *Ap* after growth in rifampin for 48 h were normalized to counts at the start of the experiment ($t$=0). Error bars are S.D. for each condition, $n$=3.

c) Inoculum size did not change the MIC of rifampin in *Lp* monocultures. Viable cell plating counts of *Lp* after growth in rifampin for 48 h were determined from different initial cell densities (numbers in colors, CFU/mL at $t$=0). Error bars are S.D. for each condition, $n$=3.

d) Inoculum size did not change the survivability of *Lp* to supraMIC concentrations of rifampin. Viable cell plating counts of *Lp* after growth in rifampin for 48 h were normalized to the initial cell density, same data as in (c). Error bars are S.D. for each condition, $n$=3.

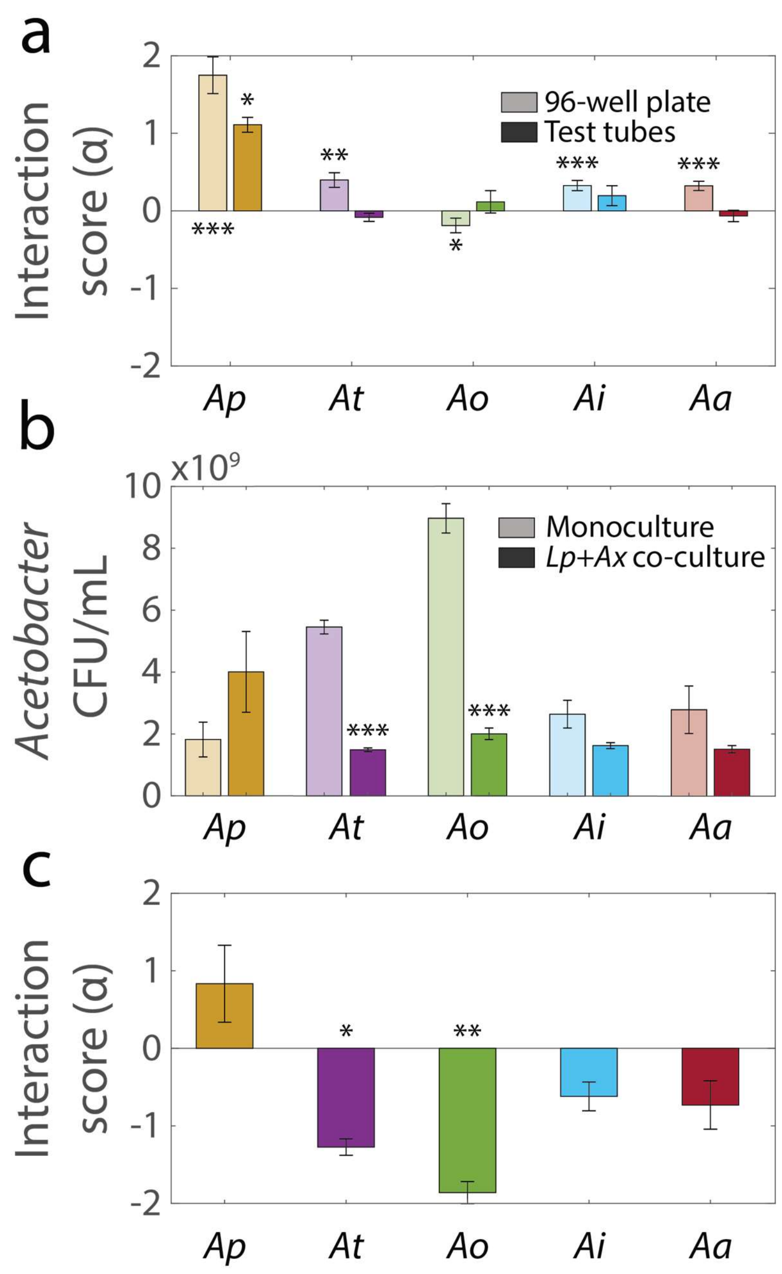


**Supplementary Figure S3. Interaction scores of *Lactobacillus plantarum* (*Lp*) and *Acetobacter* spp.**

a) Interaction score for *Lp* and *Acetobacter* spp. calculated using OD (Eq. 1 in Supplementary Text) measured after 48 h of growth in a 96-well plate (light colors) or in test tubes (dark colors). *Acetobacter orientalis*, *Ao*; *Acetobacter aceti*, *Aa*;

other species abbreviations are as defined in Figure S1. Error bars indicate standard deviation (S.D.), $n$=5 for 96-well plates, $n$=3 for test tubes. $P$-values are from a Student's two-sided $t$-test of the hypothesis that the score is 0 using the mean OD of individual samples (monocultures and co-cultures) to calculate the mean and S.D. by propagation of error (*: $P<5x10^{-3}$, **: $P<1x10^{3}$, ***: $P<1x10^{-4}$).

b) Acetobacters with *Lp* did not significantly increase *Ap* cell density, but did significantly decrease *At* and *Ao* cell densities after 48 h. Error bars are S.D. for each condition, $n$=3. $P$-values are from a Student's two-sided $t$-test of the difference from the monoculture (***: $P<2x10^{-4}$).

c) Interaction scores for *Lp* and *Acetobacter* spp. calculated from CFU/mL (Eq. 1 of Supplementary Text) measured after 48 h of growth. Error bars indicate S.D., $n$ = 3. $P$-values are from a Student's two-sided $t$-test of the hypothesis that the score is 0 using the mean OD of individual samples (monocultures and co-cultures) to calculate the mean and S.D. by propagation of error (*: $P<0.01$, **: $P<1x10^{-3}$).

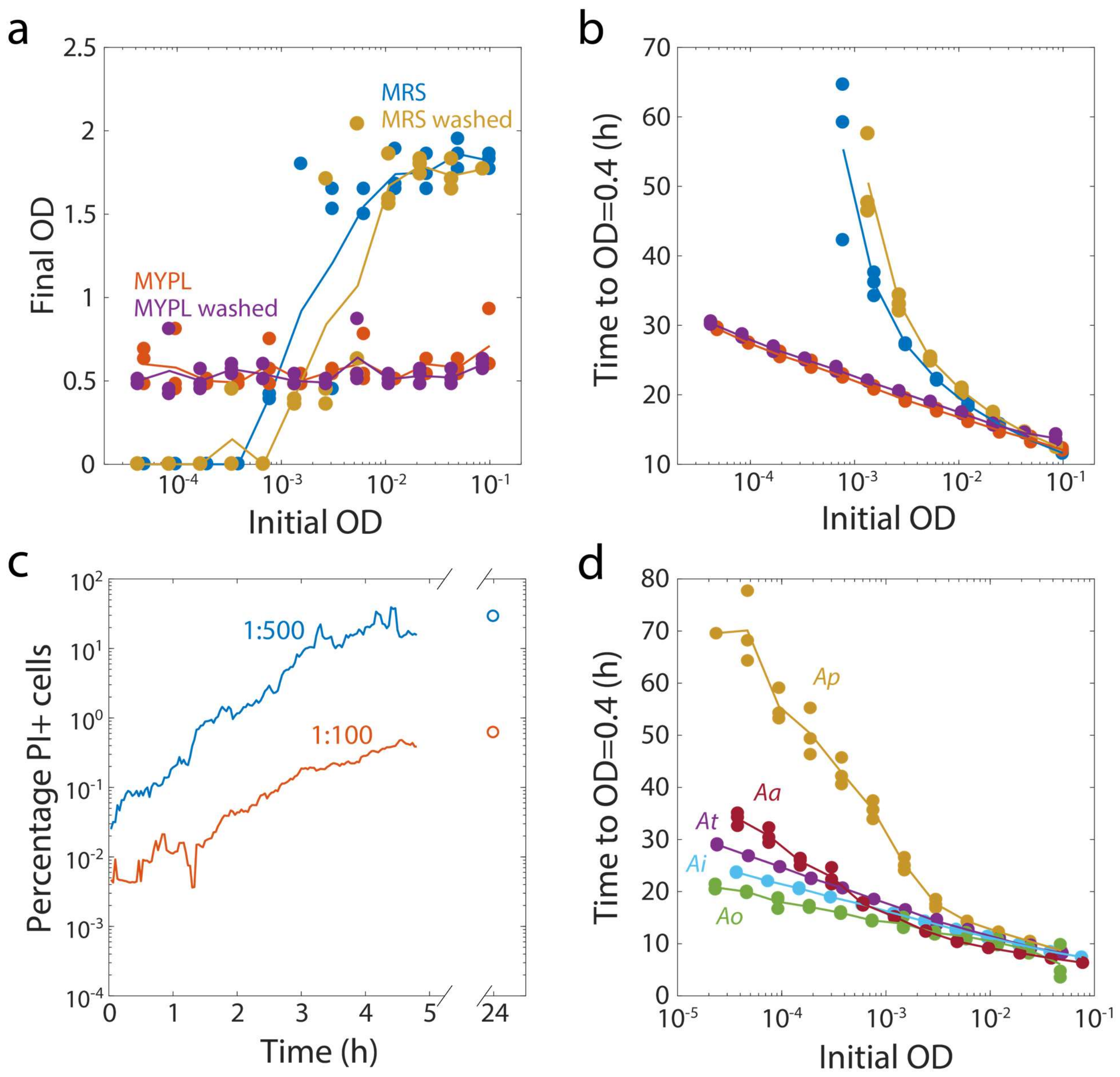


**Supplementary Figure S4: Growth of *Acetobacter pasteurianus* (*Ap*) and *Acetobacter aceti* (*Aa*) depends on inoculum size.**

a) *Ap* does not grow from low inocula in MRS but it grows independent of inoculum size in MYPL. Washing saturated cultures before diluting into fresh media to remove any metabolites accumulated does not change the inoculum-dependency of *Ap* growth. Cultures were directly diluted from a 48-h saturated culture or washed with PBS. Dots correspond to technical replicates (*n*=3) and the line is the mean OD.

b) Dependency of growth on inoculum size is evidenced by an exponential relationship between initial inoculum and time needed to reach a threshold OD. Time required to reach OD 0.4 for cultures started at various initial ODs. Dots correspond to technical replicates (*n*=3) and the line is the mean time.

c) Death of *Ap* cells in MRS is higher at low inocula. Percentage of *Ap* cells that incorporated propidium iodide (PI) over time, measured from fluorescence and phase-contrast images on an MRS pad containing PI after 100- or 500-fold dilution from stationary phase.

d) Ap and Aa growth is dependent on inoculum size. Time to reach OD 0.4 for *Acetobacter* monocultures started at various initial ODs. Dots correspond to technical replicates (*n*=3) and the line is the mean time.

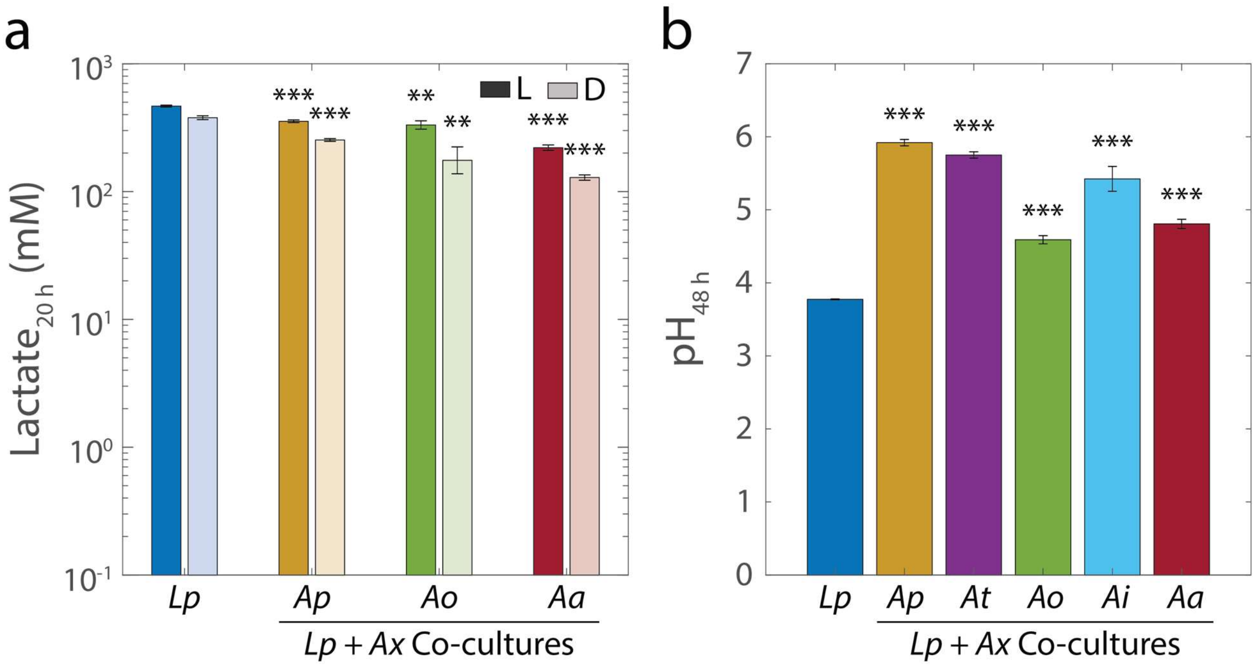


**Supplementary Figure S5. Lactate accumulates in *L. plantarum* (*Lp*) monocultures and co-cultures, and *Acetobacter*-driven consumption of lactate reverses the initial pH decrease.**

a) Lactate is accumulated in *Lp* monocultures and in co-cultures with *Ap*, *Ao* and *Aa* before entering stationary phase. L- and D-lactate concentration was measured enzymatically from the supernatants of 20-h monocultures of *Lp* or co-cultures of *Lp* with acetobacters (Methods). Species names are abbreviated as in Figure S3. Error bars are standard deviation (S.D.) for each condition, $n$=3. $P$-values are from a Student's two-sided $t$-test of the difference from the monoculture (**: $P<3\times10^{-3}$, ***: $P<3\times10^{-4}$).

b) pH in stationary phase of co-cultures of *Lp* is higher than monoculture. pH meter readings (Methods) of saturated *Lp* monocultures and co-cultures with acetobacters. Error bars are S.D. for each condition, $n$=3. $P$-values are from a Student's two-sided $t$-test of the difference from the monoculture (***: $P<2x10^{-4}$).

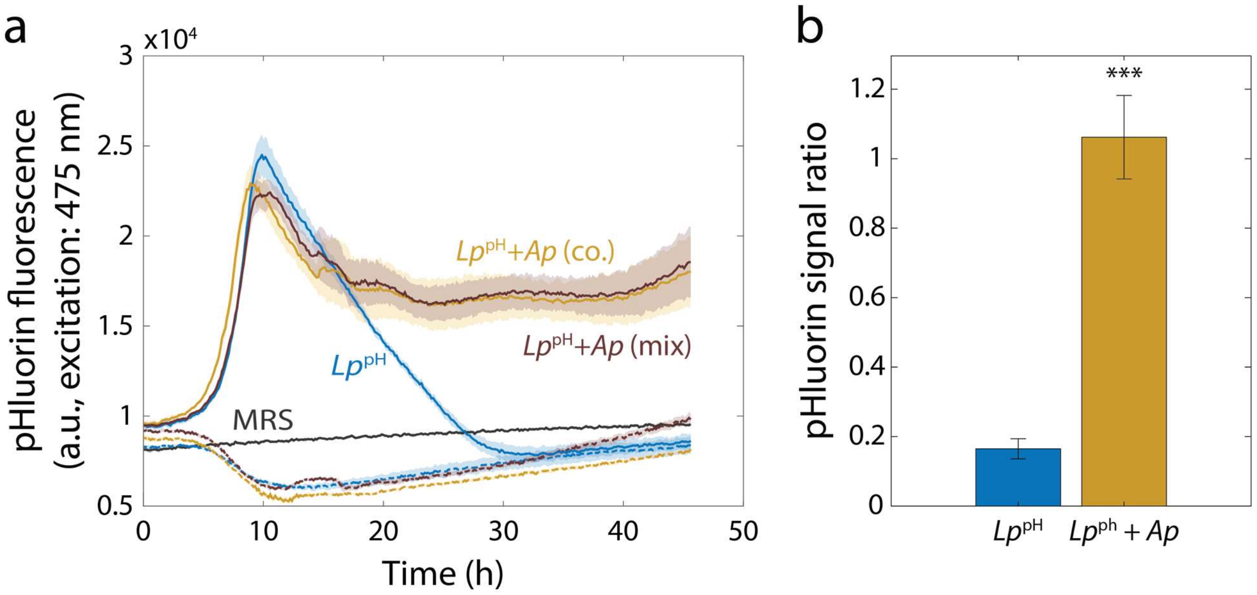


**Supplementary Figure S6: *L. plantarum* (*Lp*) intracellular pH is higher in stationary phase when grown with *Acetobacter pasteurianus* (*Ap*).**

a) Intracellular pH of *Lp* decreases upon entry to stationary phase in monoculture but not in co-culture with *Ap*. A strain of *Lp* containing a plasmid expressing the pH-sensitive GFP pHluorin ($Lp^{pH}$) was grown in monoculture, co-cultured with *Ap*, or mixed with *Ap* at the start of the experiment. Fluorescence emission (at 509 nm) was measured at an excitation wavelength of 475 nm over time, and is correlated with pH. Shaded regions indicate standard deviation (S.D.), $n$=3. Cultures of the parent strain without pHluorin (*Lp*, dashed lines) and MRS alone (gray) were included as controls.

b) Intracellular pH increases in co-cultures of $Lp^{pH}$ and *Ap* after 48 h of growth. Fluorescence (emission at 509 nm) was measured for two excitation wavelengths (405 and 475 nm) after resuspending pHluorin-harboring cells in PBS. The ratiometric signal scales with pH. Error bars are S.D., $n=3$. $P$-values are from a Student's two-sided $t$-test of the difference from the monoculture (***$P<0.001$).

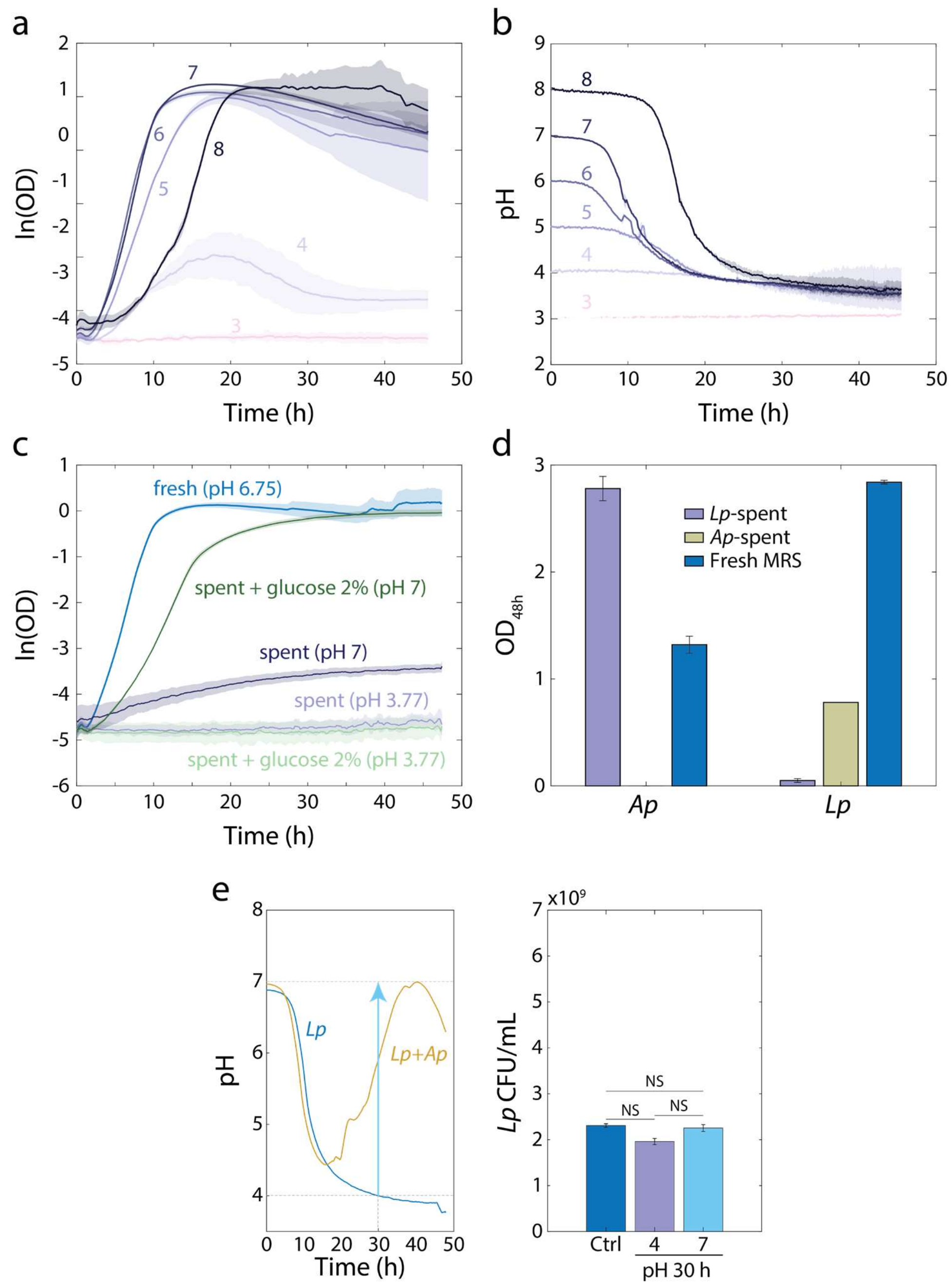


**Supplementary Figure S7: Growth of *L. plantarum* (*Lp*) in stationary phase is limited by the acidity of the medium.**

(a-b) Growth of *Lp* depends on the initial pH of the medium and it stops at a common pH of ~3.7. Logarithm of (a) OD or (b) pH (measured with the pH-sensitive dye BCECF) over time for cultures of *Lp* in MRS at various starting pH values (colors). Shaded regions are standard deviation (S.D.), $n$=4.

c) *Lp*-spent media can support growth of bulk *Lp* culture when taken from pH 3.77 to pH 7 and it supports faster growth rates and higher carrying capacity when supplemented with glucose at pH 7. Error bars are S.D., $n = 3$.

d) *Lp*-conditioned medium supports growth of *Ap* to higher OD than fresh medium and *Ap*-conditioned medium supports growth of *Lp* to lower OD than fresh medium. Error bars are S.D., $n$=3. $P$-values are from a Student's two-sided $t$-test of the difference from growth in fresh MRS (***: $P<0.001$).

e) Increasing the pH of an *Lp* monoculture at $t$ = 30 h from 4 to 7 to mimic the pH increase in *Lp*-*Ap* co-culture (left panel) does not lead to significant increase in CFU/mL with respect to a control. A 48-h-old culture with no changes in pH was used as a control (Ctrl.). Error bars are S.D. for each condition, $n$=3. $P$-values are from a Student's two-sided $t$-test of the difference between the cultures (NS, not significant, $P>0.0167$).

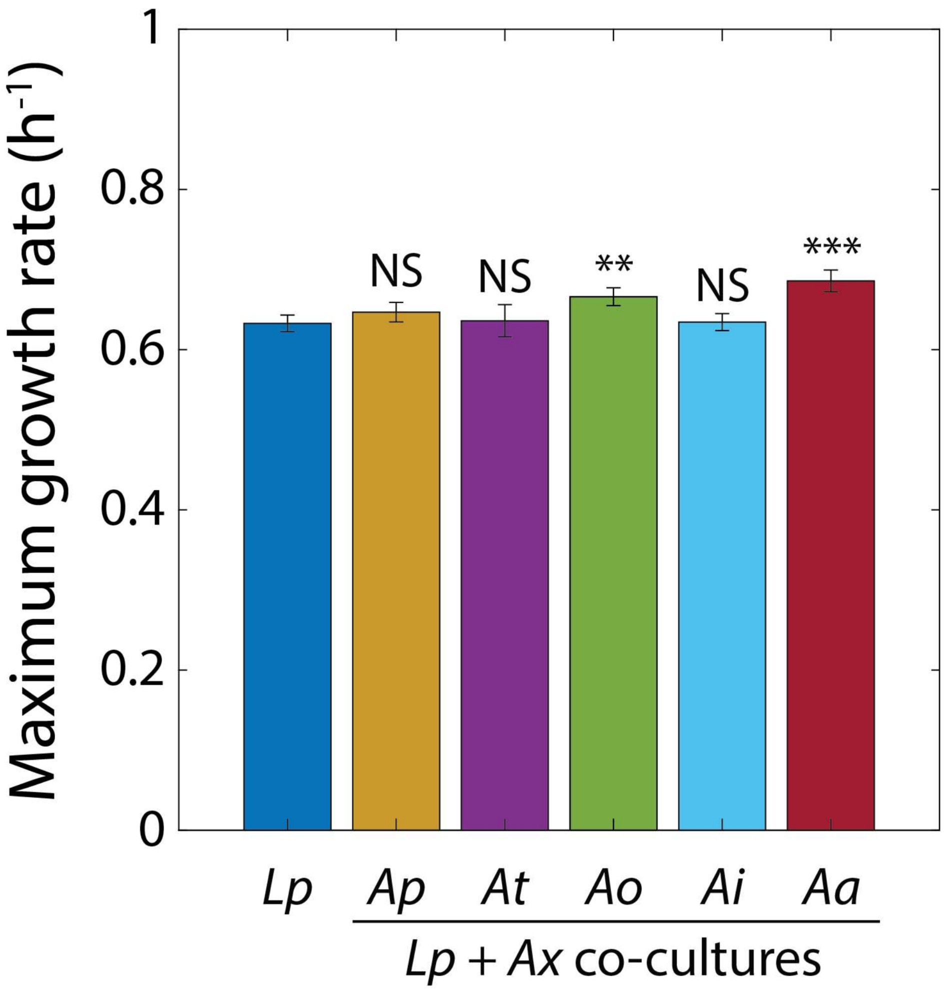


**Supplementary Figure S8: The growth rate of *L. plantarum* (*Lp*) co-cultures is similar to that of an *Lp* monoculture.** Maximum growth rate of *Lp* in monoculture or co-cultures with acetobacters (*Ax*) obtained by fitting growth curves to the Gompertz equation[1]. Species names are abbreviated as in Supplementary Figure S3. Error bars are S.D., *n*=5. *P*-values are from a Student's two-sided *t*-test of the difference from the monoculture (**: $P<2\times10^{-3}$, ***: $P<2\times10^{-4}$, NS: not significant).

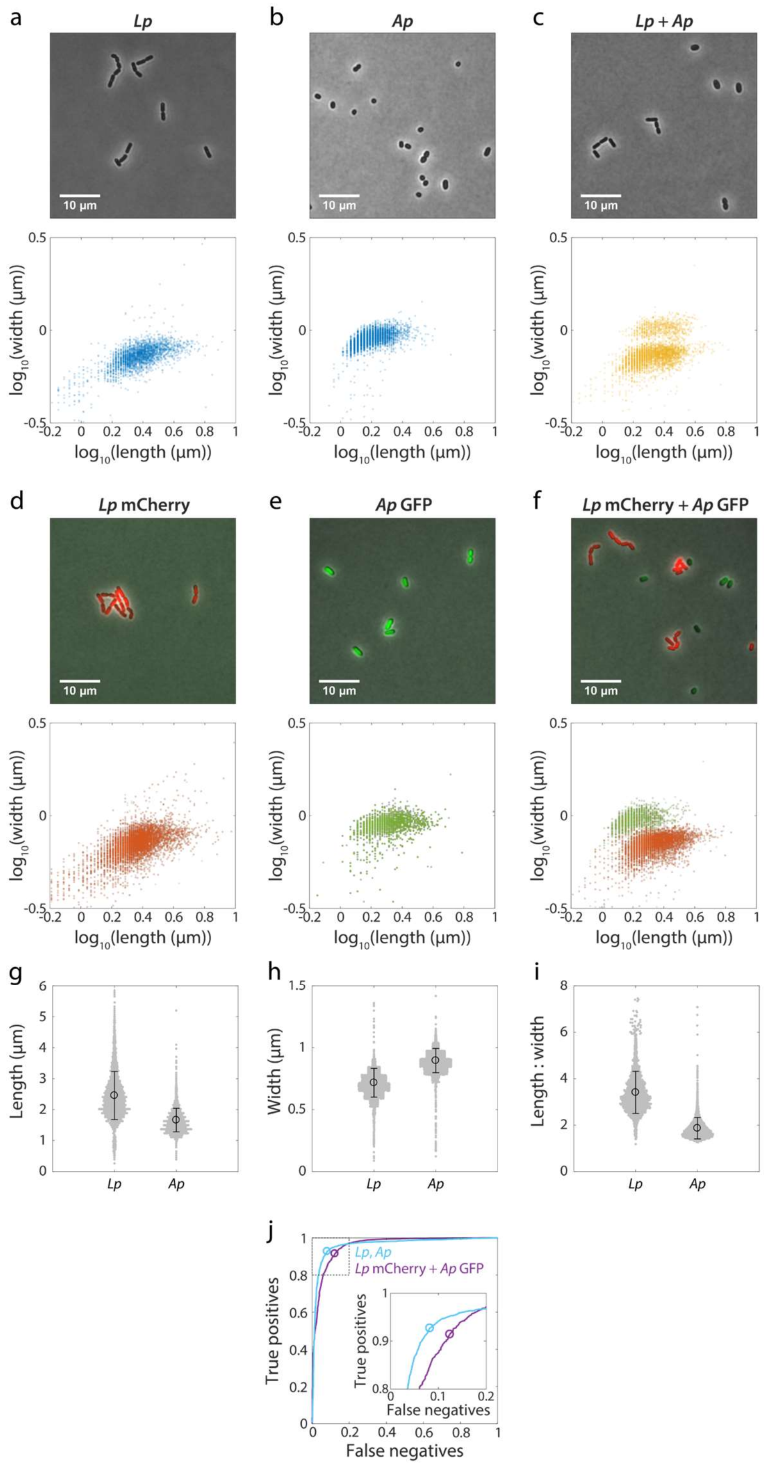


**Supplementary Figure S9: Morphological differences permit differentiation of *L. plantarum* (*Lp*) from *Acetobacter pasteurianus* (*Ap*) in phase-contrast images.**

(a-c) Representative phase-contrast images (top) enable quantification of width versus length (bottom) of individual cells in 48-h monocultures of *Lp* (a) or *Ap* (b) and in co-cultures of both species (c).

(d-f) Representative phase-contrast images are overlaid with fluorescence images (top) for quantification of width versus length (bottom) of 48-h monocultures of mCherry-tagged *Lp* (d) or GFP-tagged *Ap* (e), and co-cultures of both species (f).

(g-i) Morphology enables differentiation of Lp and Ap in phase microscopy images. Length (g), width (h), and aspect ratio (length/width, i) for monocultures in (a) and (b). Grey dots are individual cells, black circle is the mean, and error bars are standard deviation.

j) ROC curve for a classifier of *Lp* and *Ap* using the aspect ratio as a threshold. Curves shown correspond to the true-positive rate (sensitivity) of *Lp* and false-positive rate (1-specificity) of *Ap* classified as *Lp* for data in (a) and (b) (light blue) or (f) (purple). Circles show the optimal compromise points, with ln(aspect ratio) of 0.378 and 0.372 for light blue and purple, respectively. Inset: zoom-in of region inside dashed box.

### Supplementary Tables

**Supplementary Table S1: Strains used in this study.**

| Strain | Description | Colony aspect | Ref. |
|---|---|---|---|
| KC1199 | *Lactobacillus plantarum* (WF) wild fly (*D. melanogaster*) isolate | MRS: opaque white to pale yellow, circular, large, smooth surface, umbonate, smooth to irregular edge<br>MYPL: opaque white, circular, small, smooth surface, raised, smooth edge | [2] |
| KC1200 | *Lactobacillus brevis* lab fly (Oregon-R) isolate | MRS: opaque white, circular, medium size, raised, smooth edge<br>MYPL: opaque white, circular, small, raised, smooth edge. | [2] |
| KC1201 | *Acetobacter pasteurianus* lab fly (Oregon-R) isolate | MRS: opaque beige, circular, medium size, mucoid, raised, smooth edge<br>MYPL: opaque beige, circular, large, mucoid, raised, smooth edge | [3] |
| KC1202 | *Acetobacter tropicalis* lab fly (Oregon-R) isolate | MRS: opaque beige, circular, medium size, mucoid, raised, smooth edge<br>MYPL: opaque beige, circular, large, mucoid, raised, smooth edge | [3] |
| KC1203 | *Acetobacter orientalis* lab fly (Oregon-R) isolate | MRS: opaque beige, irregular, medium to large size, dry, flat to irregular surface, smooth to lobate edge<br>MYPL: opaque beige, irregular, medium to large size, dry, flat to irregular surface, smooth to lobate edge | [2] |
| KC1204 | *Acetobacter indonesiensis*s lab fly isolate | MRS: opaque beige, circular, medium size, mucoid, raised, smooth edge | [2] |

| | | | |
|---|---|---|---|
| | | MYPL: opaque beige, circular, large, mucoid, raised, smooth edge | |
| KC1205 | *Acetobacter aceti* lab fly isolate | MRS: opaque beige, circular, medium size, mucoid, raised, smooth edge<br>MYPL: opaque beige, circular, large, mucoid, raised, smooth edge | [2] |
| KC1206 | *Lactobacillus plantarum* (WF) wild fly isolate pCD256-*P*11-mCherry | | [2] |
| KC1207 | *Lactobacillus plantarum* (WF) wild fly isolate pCD256-P11-pHluorin | | This work |
| KC1208 | *Acetobacter pasteurianus* lab fly (Oregon-R) isolate pCM62-*P*lac-sfGFP | | This work |

**Supplementary Table S2: Minimum inhibitory concentrations of antibiotics for the four primary members of the fruit fly gut microbiota.** Minimum inhibitory concentrations are expressed in μg/mL ± standard deviation, *n*=3.

| **Antibiotic** | **Class** | ***L. plantarum*** | ***L. brevis*** | ***A. pasteurianus*** | ***A. tropicalis*** |
|---|---|---|---|---|---|
| Ampicillin | Beta-lactam | 10.42±3.61 | 25±0 | 16.67±7.22 | >100 |
| Streptomycin | Aminoglycoside | >50 | 50±0 | 12.5±10.83 | 1.56±0 |
| Chloramphenicol | N/A | 7.29±4.77 | 6.25±0 | >50 | 25±0 |
| Tetracycline | N/A | >50 | >50 | >50 | 1.56±0 |
| Erythromycin | Macrolide | 0.2±0 | 0.2±0 | >25 | >25 |
| Ciprofloxacin | Fluoroquinolone | >50 | 25±0 | >50 | 12.5±0 |
| Trimethoprim | N/A | >50 | >50 | >50 | >50 |
| Spectinomycin | N/A | 200±0 | 200±0 | 66.67±28.87 | 25±0 |
| Rifampin | N/A | 1.3±0.45 | 0.52±0.23 | 6.25±0 | 4.17±1.8 |
| Vancomycin | N/A | >100 | >100 | >100 | >100 |
